# Inferring cellular communication through mapping cells in space using Tangram 2

**DOI:** 10.1101/2025.09.28.679077

**Authors:** Hejin Huang, Alma Andersson, Sunny Z. Wu, Gabriele Scalia, Jenna Collier, Kam Hon Hoi, Sören Müller, Graham Heimberg, Hector Corrada-Bravo, Shreya Gaddam, Jan-Christian Hütter, Shannon Turley, David Richmond, Aicha BenTaieb, Tommaso Biancalani

**Author notes:** these authors contributed equally to this work (Hejin Huang, Alma Andersson).

## Abstract

Cell-to-cell communication (CCC) shapes development, immunity, and disease, yet current spatial transcriptomics (SRT) platforms rarely achieve both single-cell resolution and high-quality transcriptome-wide coverage to accurately characterize CCC in the tissue microenvironment. Tangram2 bridges this gap by integrating single-cell RNA sequencing (scRNA-seq) with SRT to identify genes whose expression changes as a function of neighboring cell types. By accurately mapping cells in tissue space, Tangram2 disentangles interaction-driven transcriptional shifts from intrinsic identity markers, yielding mechanistic CCC maps. Validation across diverse settings—including Slide-tags, co-cultured experiments, human lymph nodes, and simulations—demonstrates high accuracy. Applied to triple-negative breast cancer (TNBC) and cutaneous squamous cell carcinoma (cSCC), Tangram2 recapitulates known biology and uncovers new hypotheses, including various immunosuppressive mechanisms in TNBC and a macrophage–regulatory T-cell circuit associated with survival in cSCC.

## INTRODUCTION

Cell-to-cell communication (CCC) underlies development, tissue homeostasis, immune defense, and regeneration. Its disruption can directly drive the pathogenesis of major disease, including cancer, cardiovascular and neurodegenerative disorders, and various forms of impaired tissue repair. Consequently, CCC is a critical focus for biomarker discovery and development of precision therapeutics ^1–3^. Early efforts to chart CCC relied on low-throughput, hypothesis driven assays for specific ligand-receptor interactions ^4 5^. Over the past two decades, transcriptomic profiling, particularly single-cell RNA sequencing (scRNA-seq), has transformed the field by enabling transcriptome-wide resolution of cellular states, while expert-curated LR resources compiled from the literature and public databases provided the priors needed for systematic CCC analysis ^6,7^. In parallel, computational methods have been developed to infer cellular crosstalk from transcriptomic data, first using bulk or scRNA-seq ^7–10^, later using spatially resolved transcriptomics (SRT) ^11–14^, and more recently with integrative approaches that combines multiple modalities and resources ^15^.

Transcriptomics-based computational methods for CCC inference typically combine ligand and receptor expression with prior knowledge to score ligand–receptor interactions from ‘sender’ to ‘receiver’ cell types ^7,8^ or to predict ligand-to-target gene program, with additional distance/contact constraints in spatial methods ^16–18^. However, three challenges limit their effectiveness. First, identifying interacting subpopulations remains nontrivial: scRNA-seq lacks spatial context, while most SRT platforms either lack true single-cell resolution or sacrifice transcriptome-wide coverage. Second, differential expression is confounded by intrinsic factors, such as cell-type identity programs which may obscure interaction-specific signals. Third, validation remains difficult due to the scarcity of ground-truth datasets, leaving most methods supported by qualitative case studies rather than systematic benchmarking.

We introduce Tangram2, a computational method for inferring CCC that addresses the challenges outlined above. Tangram2 comprises three modules (**Figure 1**). Tangram2-mapping generates spatial probability maps for scRNA-seq profiles by integrating matched SRT data, enabling identification of co-localized cell populations without sacrificing gene coverage. Mapping scRNA-seq in space was pioneered in embryonic systems ^19^ and later extended to chart common coordinate frameworks of human organs ^20^, with recent advances highlighting developmental trajectories ^21^. Tangram2-CCC applies a linear model to the mapped data to infer gene-level communication coefficients, identifying interacting populations while disentangling intrinsic cell-type signals from interaction-driven programs. Finally, Tangram2-evalkit provides a framework for generating synthetic data seeded from real datasets, which we used for systematic benchmarking. Tangram2 builds on prior approaches ^22–26^ but places stronger emphasis on modeling spatial cell–cell interactions.

**Figure 1:**
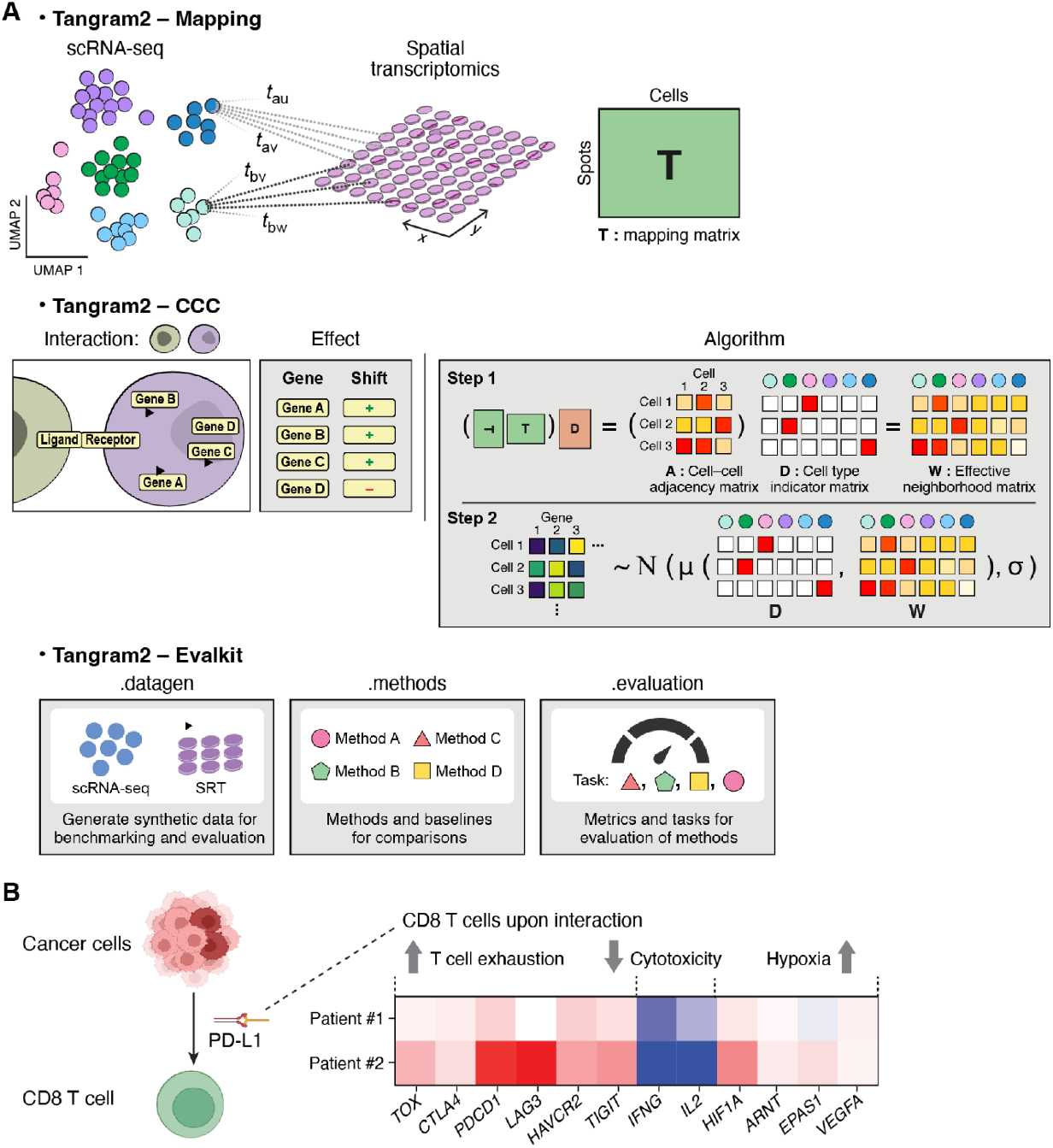
Schematic Diagram of Tangram2. **A**. Tangram2 comprises three modules **Tangram2-mapping** aligns the single-cell RNA-seq to spatial transcriptomics. **Tangram2-CCC** investigates effects of cell-cell interaction based on the cell adjacency information. **Tangram2-evalkit** creates a workflow for synthetic data generation, method evaluation and benchmarking. **B**. An example of cell-cell interaction effects captured in tumor microenvironment of TNBC: T cell exhaustion and hypoxia markers are upregulated while the cytotoxic markers are downregulated in CD8 T cells upon interactions with PD-L1+ cancer

We validated Tangram2 across multiple settings (**Figure 2**). First, we assessed Tangram2-mapping using an established benchmarking pipeline ^27^ as well as synthetic datasets generated by Tangram2-evalkit for mapping and deconvolution, demonstrating its state-of-the-art performance. Next, we evaluated Tangram2-CCC in four settings, each combining scRNA-seq and SRT data, but with ground-truth cell–cell interaction information derived in distinct ways: (i) Slide-tag data synthetically down-sampled to Visium-like resolution, used to predict interaction-induced genes ^28^; (ii) an in vitro co-culture of dendritic cells and T cells, serving as ground truth for interactions between these cell types ^29^; (iii) helper T cells and germinal center B cells from annotated from Gene Ontology ^30,31^; and (iv) simulations with known parameters, enabling quantitative performance assessment. Across all settings, Tangram2 reliably recovered ground-truth CCC with significant accuracy.

**Figure 2:**
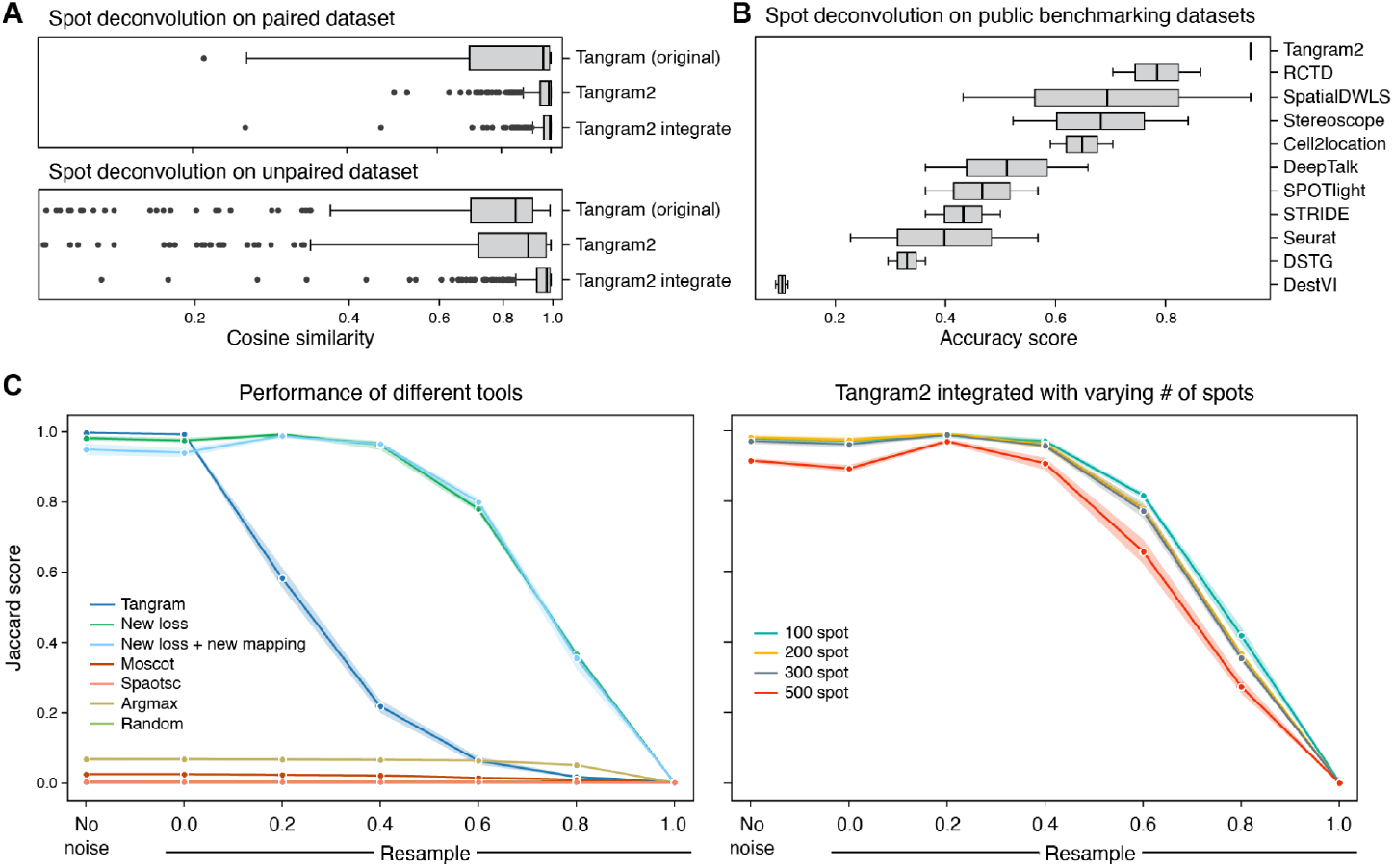
Tangram 2 demonstrates state-of-the-art integration of transcriptomics data. **A**. Comparison of Tangram and Tangram2 on spot deconvolution task based on synthetic paired and unpaired datasets from Tangram2-evalkit **B**. Benchmarking Tangram2 against all publicly available tools based on the pseudobulked SRT datasets from Li et al, in which accuracy score is based on the ranking of four metrics, including cosine similarity, Pearson coefficient, RMSE and JS divergence. **C**. Evaluation of mapping accuracy, the left panel is the comparison of Jaccard scores between all mapping tools at different noise levels when the total number of spots is 100, while the right panel shows the performance of Tangram2 with respect to the varying number of spots from 100 to 500.

As a case study, we applied Tangram2 to investigate interactions between immune populations and malignant cells within the tumor microenvironment (**Figure 4** and **5**). We focused on two cancer types: triple-negative breast cancer (TNBC) and cutaneous squamous-cell carcinoma (cSCC), both of which are characterized by immunosuppressive microenvironments that hinder effective anti-tumor responses ^32–34^. In both cases, Tangram2 successfully recapitulated established biological mechanisms while revealing novel hypotheses. In TNBC, Tangram2 identified several immunosuppressive mechanisms, including: PD-L1-induced T-cell exhaustion in the tumor microenvironment, CAF-driven tumor growth, as well as macrophage and T-reg induced immune suppression of CD8^+^ T cells. In cSCC, Tangram2 identified a macrophage–Treg circuit positively associated with longer survival and an association between TSK composition and immunosuppression in the tumor microenvironment.

**Figure 3:**
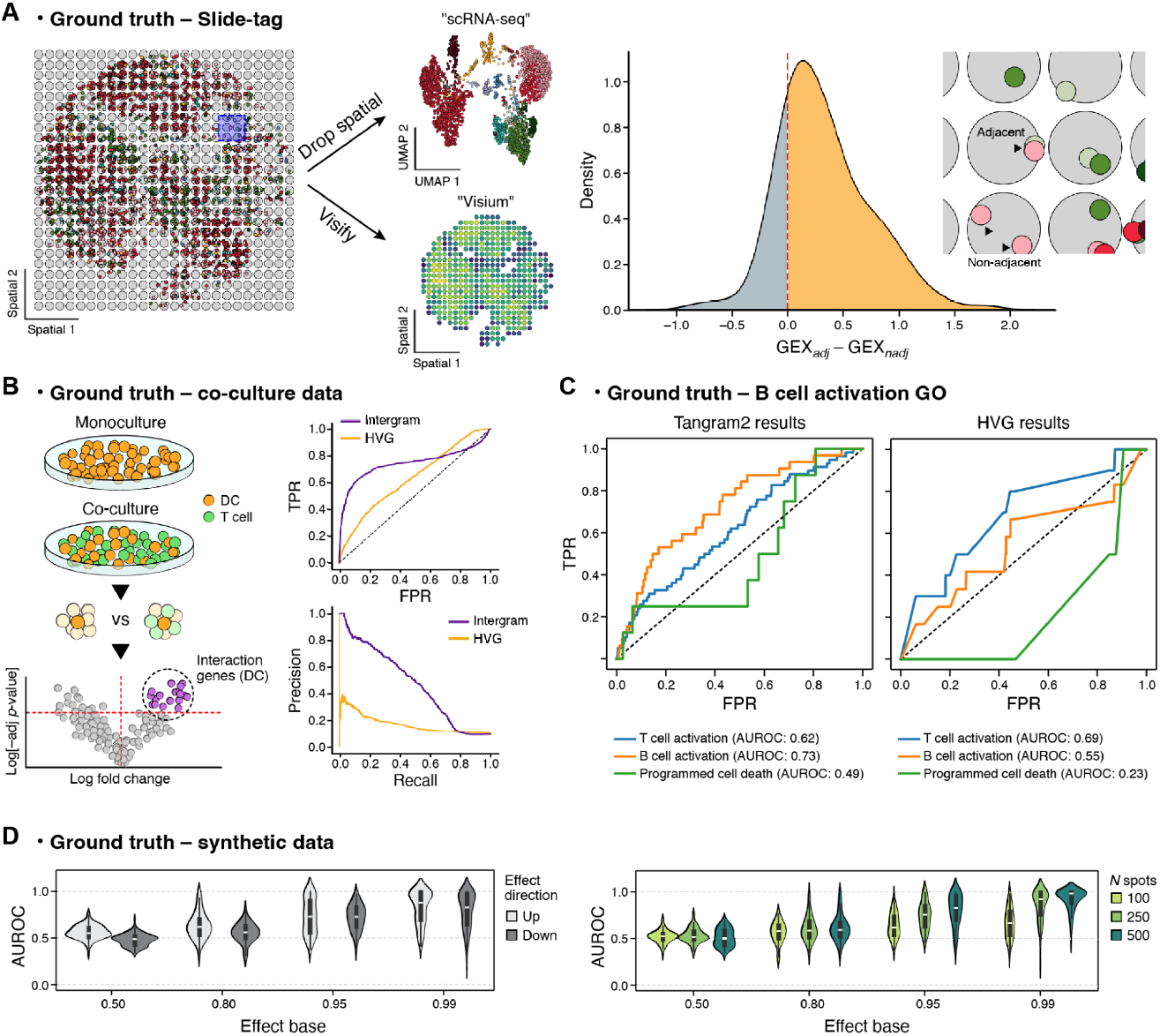
Tangram2 is validated by accurately identifying cell-cell interaction genes in semi-synthetic and experimental datasets. **A**. Tangram2 results on semi-synthetic data from Slide-tag: The distribution of mean expression differences between adjacent (adj) and non-adjacent (nadj) cells for the top ten genes (ranked by interaction coefficients) across cell-type interactions. Yellow indicates positive differences (desirable), gray indicates negative differences. The inset zooms into an area in panel A, showing examples of adjacent and non-adjacent cells for the interaction B_naive ← NK (pink ← light green). **B**. Tangram2 performance on the mouse colon dataset compared to cell co-culture experiment: we compare DCs in monocultures and co-cultures with T-cells to identify differentially expressed genes, used as ground truth interaction genes for the interaction DCs ← T-cells. **C**. Results from the B cell activation analysis: the figure shows the receiver operating characteristics (ROC) curve for Tangram2-CCC and genes ranked by dispersion from a highly variable gene (HVG) analysis. B-cell activation should have the highest AUC (area under curve), as the other two pathways are negative controls. **D**. AUROC of interaction predictions performance on the synthetic data with artificial interactions when stratified by two different axes effect_base (baseline expression) and either of effect_direction and n_spots.

**Figure 4:**
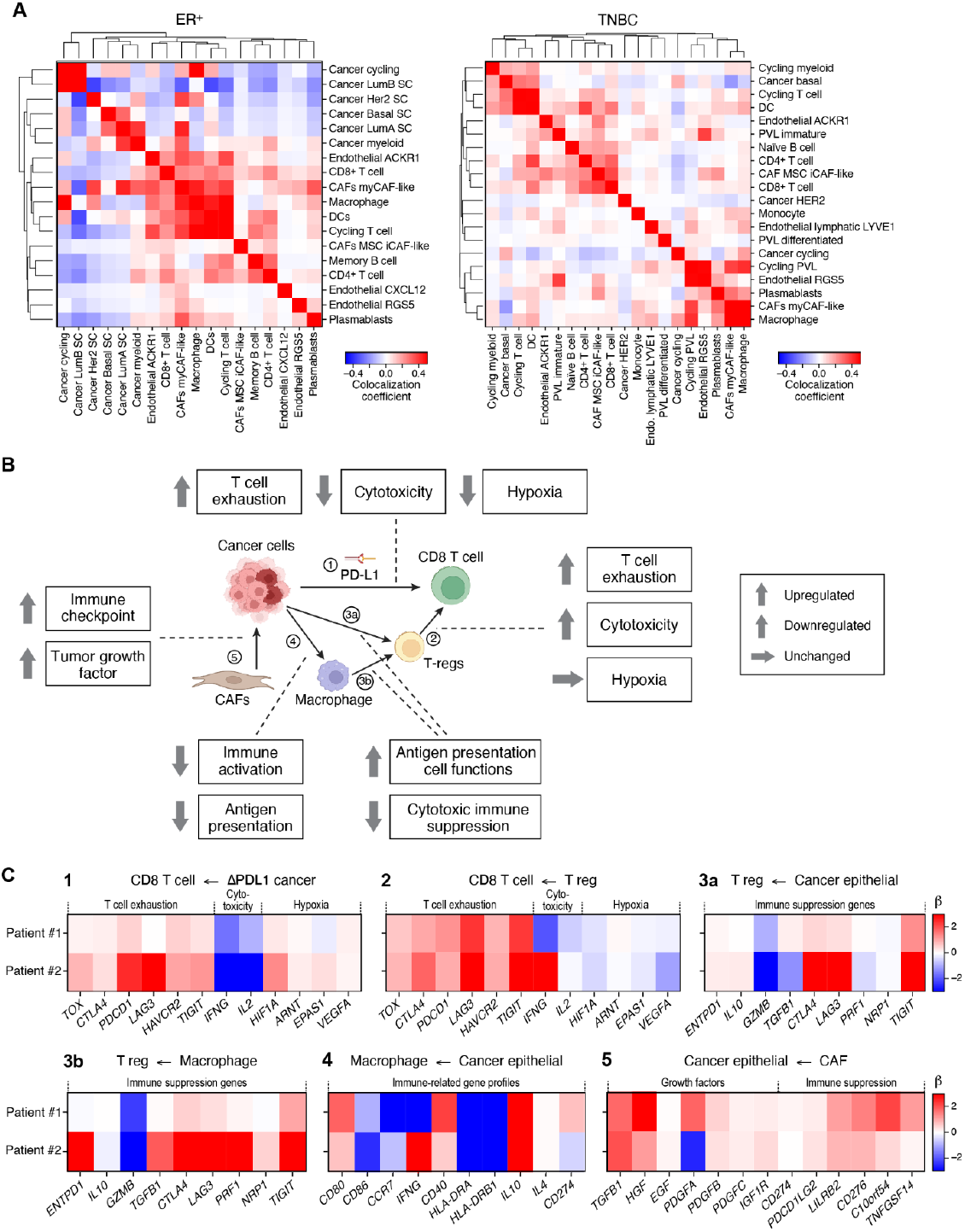
Tangram2 identifies immunosuppressive paths between cancer and T cells in TNBC. **A**. Clustermap of the average colocalization scores between cell types of TNBC cancer samples based on the Tangram2-mapping results of two breast cancer samples. The cell types that consist of less than 0.05% of total cell populations in the spatial samples are excluded from this analysis **B**. Summary of the direct and indirect immunosuppressive paths in the tumor microenvironment of TNBC discovered by Tangram2-CCC **C**. The interaction coefficients of selected gene lists learned by Tangram2-CCC between communicating cell types highlighted in B. For each interaction, the list of genes is selected based on the affected cell type. For example, when CD8 T cell is the affected cell type (in both *CD8 T cell <– PDL1 cancer* and *CD8 T cell <– T-regs*), the selected gene list consists of markers for T cell exhaustion, T cell cytotoxicity and hypoxia.

## RESULTS

### Tangram2-evalkit generates synthetic data for mapping and cell-cell communication evaluation

We start with a scRNA-seq dataset with annotated cell types, which serves as the seed for synthetic data generation (**Supplementary Figure 1A**). From this, we construct a square grid of spots, similar to the layout of capture-based SRT technologies such as 10x Visium. Each spot is populated with cells sampled from the seed scRNA-seq dataset, with a pre-specified average number of cells and cell types across all spots, however these numbers differ between individual spots. This setup allows us to mimic different assay and tissue properties (*e*.*g*., high vs. low spatial resolution and homo- vs. heterogeneous tissues). We also provide the option to impose spatial coherence in the arrangement of spots with respect to cell type distribution.

We evaluate model robustness and sensitivity by resampling transcripts in individual cells with and without multinomial noise, simulating varying levels of biological and technical variability.^35,36^. We then add synthetic intercellular interactions (**Methods**). Each interaction involves a single “signal” gene, and a set of downstream “effect” genes. For any given interaction, cells are assigned to one of three mutually exclusive classes: signalers, receptive cells, or non-receptive cells. Signaler cells overexpress the signal gene and can activate nearby receptive cells; non-receptive cells remain unaffected. All cells may express both signal and effect genes, but expression levels are dependent on whether they are a signaling cell, a receptive cell actively receiving a signal, or a non-interacting cell (**Methods**). The assignment of signalers and receptive cells can be constrained to specific cell types or applied broadly. Importantly, the addition of effect genes does not alter the underlying cell identity (**Supplementary Figure 9**).

To preserve spatial structure, we designate spots as either active or inactive based on a fixed probability (default: 0.5). Signalers are located in active spots. Receptive cells are then activated with a probability parameter, provided they co-localize with signalers. Synthetic expression levels for both signal and effect genes are then introduced accordingly.

The final output of this process is a synthetic paired single cell and spatial dataset containing interacting cells and their downstream effects. The corresponding single-cell data consists of all gene expression profiles from the cells assigned to a spot in the synthetic spatial data, with their added signal and effects genes.

### State-of-the-art mapping using Tangram2-mapping

Mapping refers to the process of assigning individual cells from scRNA-seq data, denoted as 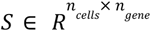 to spatial coordinates in SRT data denoted as 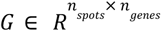. We call datasets paired, when scRNA-seq and SRT data originates from the same tissue section (ideally from adjacent slices), and unpaired for distinct but biologically similar samples, such as tissues from different patients with the same tumor type. Ideally, mapping methods should perform robustly across both settings.

Tangram2-mapping, like Tangram and its extension ^26,37^, tackles this integration challenge by casting it as an optimization problem. The method works as follows (**Methods**): it first identifies a set of training genes, typically consisting of the marker genes from each cell type; next, the algorithm learns the mapping matrix 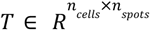 that gives the mapping probability from cells to spatial spots by ensuring that the expression of the training genes aligns with that of the SRT data. Tangram2-mapping incorporates two key innovations. Firstly, one of the key components of the loss function characterizing alignment has been changed as below (**Methods**):

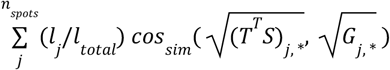

Here 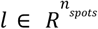 denotes the number of transcriptomes measured at each spot. The Bhattacharyya distance (square root of cosine similarity) improves the mapping of sparse gene expression, while the *l*-weighted spot-level aggregation aims to make the alignment less susceptible to the selection of low-quality training genes ^38^. Second, we include a new mapping mode, named integrated mapping, specifically crafted to deal with unpaired datasets. This new mode first estimates the cell type proportions of SRT, and then renormalizes the scRNA-seq with the proportion before performing standard mapping, helping to correct the mismatch between scRNA-seq and SRT thus enhancing the mapping accuracy (**Methods)**.

We first benchmarked Tangram *vs*. Tangram2-mapping using synthetic datasets generated using Tangram2-evalkit seeded from breast cancer dataset^39^ which included both paired (same-patient) and unpaired (cross-patient) configurations. Both Tangram2-mapping modes outperformed original Tangram on paired data (cosine similarity >0.95), but only the Tangram2 integrated mapping approach maintained high accuracy on unpaired data(**Figure 2A**). Furthermore, ablation study on the new loss function shows that the Bhattacharyya distance improves the overall alignment score, while the spot-level aggregation improves the mapping robustness when low-quality genes are included (**Supplementary Figure 2**).

Next, we benchmarked Tangram2-mapping against ten public deconvolution tools: Cell2Location^40^, RCTD^41^, SpatialDWLS^42^, Stereoscope^43^, SPOTlight^44^, STRIDE^45^, Seurat^46^, DeepTalk^47^, DSTG^48^ and DestVI^49^. Note that most of these tools perform deconvolution without mapping cells, ie, they estimate cell type composition within each SRT spot but do not retrieve explicitly the spatial coordinates of the single cell profiles. Using an established benchmarking protocol in which all ten tools had been previously evaluated, Tangram2-mapping achieved state-of-the-art performance (Figure 2B and Supplementary Figure 3).

To evaluate mapping, we generated synthetic SRT by Tangram2-evalkit from the scRNA-seq of cutaneous squamous cell carcinoma^50^, yielding paired data with known cell locations. The Jaccard score between the predicted position of each cell and its true position is used to evaluate the alignment of the mappings. To further contextualize the performance of the mapping methods two baseline approaches were included in the evaluation applied (**Methods**): random spot assignment (‘random’ baseline) and assignment of cells to the spot with the highest Pearson correlation (‘argmax’ baseline). Different levels of noise are introduced as indicated in the x axis, to create ‘mismatch’ between single cell and SRT.

We benchmarked four tools that perform cell-level mapping: Tangram, Tangram2-mapping (cell-level and integrated modes), and SpaOTsc ^5121^. As shown in the left panel of **Figure 2C** (100 spots), Tangram and Tangram2-mapping substantially outperform all other methods. SpaOTsc performs only slightly better than random, and worse than the argmax baseline. All Tangram-based methods achieve high accuracy at low noise levels; as noise increases, Tangram2-mapping remains more robust than Tangram, maintaining a Jaccard score ∼0.8 at noise level 0.6. Extended results with 200, 300, and 500 spots (Fig. S4) confirm these trends: performance declines as spot count increases, but Tangram2-mapping remains robust (**Figure 2C** right panel). Since the SRT is generated by paired scNRA-seq, Tangram2-mapping’s cell-level and integrated modes perform similarly, as the cell type compositions of the two datasets are equivalent (**Supplementary Figure 1B**).

### Identifying cellular communication effects via Tangram2-CCC

Tangram2-CCC is the module designed to identify cell-cell communication effects after mapping scRNA-seq to SRT data. By cell-cell communication effects, we refer to the downstream transcriptional changes in a “receiver” cell type (denoted *k*) that result from signaling by a “sender” cell type (denoted *l*), where *k* and *l* are not necessarily distinct. For example, in the case of ligand-receptor signaling, cells of type *l* express the ligand, and cells of type *k* express the corresponding receptor (*k←l*). Tangram2-CCC quantifies the effect of such interactions via the interaction coefficients β_*klg*_ which represent the effect of cell type *l* on the expression of gene *g* in cell type *k*.

Strictly speaking, Tangram2-CCC does not require SRT data as direct input. Instead, the required input is any mapping matrix 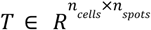 that relates cells to spots. In our analysis, we use maps from Tangram2-mapping since it showed superior performance to other methods. From that matrix, a cell-cell adjacency matrix can be derived via *A* = *TT*^*T*^. Indeed, a key insight of Tangram2, is that leveraging this mapping matrix to derive the adjacency matrix allows us to retain cell-level rather than spot-level insights.

Next, we combine the cell type indicator matrix 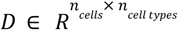 (*D*_*ik*_ = 1 if cell i is of cell type k, zero otherwise), obtained from the cell-level annotations, with the cell-cell adjacency matrix to produce the most central element to our method: the effective neighborhood matrix 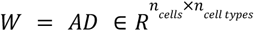. This matrix represents the effective neighborhood, or niche, of each individual cell, that is: *W*_*ik*_ reflects the abundance of cell type k in the spatial neighborhood of cell *i*. The *D* and *W* matrices are essential, because they allow us to model the single-cell gene expression as a function of each cell’s identity (cell type) and its environment (effective neighborhood). More precisely, if *X* denotes the normalized scRNA-seq data (**Methods**), we assume that this the expression of gene g in cell i can be modelled with a normal distribution: *X*_*ig*_ ∼ *N*(μ_*ig*_, σ _*g*_), with 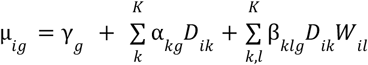. The three parameters (γ, α, β) can be explained as follows: (1) baseline expression for each gene γ, (2) an expression component specific to cell types, parametrized by 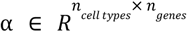, and (3) an environmental contribution shaped by the surrounding cell types which depends on 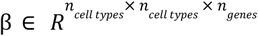. The key parameter of interest, for cell-cell interaction, is β which tells us how a gene is impacted in cell type A when in proximity to cell type B, we will refer to this β as the “interaction tensor” and the individual values as “interaction coefficients”

### Tangram2-CCC predicts cell-cell communication effects in co-localized cells

In our first validation, we assessed Tangram2-CCC’s ability to correctly infer the downstream effects of cell-cell interactions performance. For this purpose, we used a human tonsil Slide-tags dataset ^28^. Slide-tags provides spatially encoded single-nucleus RNA-seq profiles although for consistency, we will refer to nuclei as “cells”. From the Slide-tags data, we constructed paired coarse SRT and scRNA-seq datasets, integrated them with Tangram2-mapping, then inferred the cell–cell interaction tensor using Tangram2-CCC. We benchmarked the predictions of the model against the original Slide-tags measurements.

Specifically, we coarse-grained the Slide-tags data by overlaying a grid and treating each node as a mock Visium spot (**Figure 3A**). Cells within a set distance were assigned to the nearest spot; the rest was excluded. For each spot, gene expression was computed as the sum of transcripts from its assigned cells, generating “Visified” spatial data. In parallel, we removed the spatial coordinates from the Slide-tags cells, yielding an scRNA-seq-like dataset. We then applied Tangram2-mapping to map this scRNA-seq-like dataset onto the Visified spatial data, using the resulting mapping matrix as input to Tangram2-CCC.

If Tangram2-CCC worked as intended, genes with positive interaction values in the interaction tensor β should be upregulated in those receiver cells *located near* the signaling cells. To evaluate if this held true, we tested whether the top 10 genes of each interaction, ranked by their interaction coefficient, had higher expression in the receiver cells when near one or more signaling cells in the Slide-tags data. Proximity was determined using the original cell coordinates prior to Visification, applying a distance threshold smaller than that used for coarse-graining. Notably, for 1261 of 1560 genes (80% of genes; p-value < 0.01, binomial test), gene expression was increased in the adjacent cells attesting to the method’s ability to retrieve true signals (**Figure 3A**).

### Tangram2-CCC predicts cell-cell communication effects between dendritic cells and T-cells

For the next validation, we analyzed scRNA-seq data from mono- and co-cultures of splenic dendritic cells (DCs) from mice, together with CD4^+^ T cells ^29^ (**Methods**). These two cell types are known to interact, as DCs initiate and modulate T cell responses by presenting antigens and providing co-stimulatory signals. Therefore differential gene expression between a DC mono-culture, and those co-cultured with T-cells would reveal genes upregulated as a result of DC ← T cell interactions. We considered genes with an adjusted p-value ≤ 0.05 and positive log fold change as differentially expressed (**Methods)**. This yielded 1317 differential genes (Wilcoxon rank-sum; p-value ≤ 0.05; **Methods**). that we used as ground truth.

We then analyzed a second dataset from mouse colon scRNA-seq and paired Visium spatial transcriptomics, which also included annotated DCs and T-cells ^52^. Following the co-culture experiment, we focused on the interaction coefficients between DCs (receivers) and CD4+ T-cells (signaling). Of the 1317 “ground truth” genes obtained from the co-culture data, 657 (50%) overlapped with the top 1317 genes ranked by interaction coefficients from Tangram2-CCC (p-value < 0.01, hypergeometric test). To compute AUROC and AUPRC, we intersected the gene sets from the co-culture and colon scRNA-seq datasets, treating ground truth (GT) genes as positives and all others as negatives. Genes were then ranked by interaction coefficient (log fold change), and the resulting AUROC for Tangram2-based ranking was 0.74 (**Figure 3B**). We compared this result to a baseline based on the top 1317 highly variable genes in dendritic cells (DCs). Only 286 ground truth genes overlapped with the top highly variable genes, yielding an AUROC of 0.63 (i.e. 0.11 lower than the AUROC achieved by Tangram2; **Fig. 3B**).

### Tangram2-CCC predicts B-cell activation genes

For our third validation, we investigated whether Tangram2 could recover gene expression patterns associated with T cell–dependent B cell activation, which is a well-characterized and extensively studied interaction ^53,54^. For this, we used scRNA-seq and Visium data from the human lymph node ^55^, a tissue where immune cells undergo activation, differentiation, and proliferation, contributing to immune responses. We specifically focused on the interaction between CD4+ follicular T-helper cells, and germinal center B-cells that have not yet differentiated into long-lived plasma cells and memory B cells (interaction: B_naive ← T_CD4+_Tfh).

In this case, we constructed a ground truth by using publicly available gene sets that annotate different processes, including those associated with cell-interactions and compared it to Tangram2’s predictions. Specifically, we selected three Gene Ontology (GO) pathways (**Methods**): B cell activation (as ground truth), and T cell activation and programmed cell death as negative controls. We then calculated the AUROC score for all three pathways, using the absolute values of the interaction coefficients (since it is not known *a priori* if the genes are up-or downregulated) as our ranking of the genes. The scores were as follows: B-cell activation, 0.75, T-cell activation 0.64, and programmed cell death 0.56. When comparing this with a rank based on the variability of the genes within the B_naive cell types, the rankings were: B-cell activation – 0.55, T-cell activation – 0.69, and programmed cell death – 0.23, see **Figure 3C**. In short, Tangram2-CCC correctly showed the strongest activation for B-cell activation, and much lower signal for the T-cell activation, while the highly variable gene approach incorrectly flipped the order of these.

### Tangram2-CCC achieves high accuracy on synthetic data

In earlier validations, we reported performance on real datasets, although without direct validation measurements. For our final validation, we used the synthetic data to assess Tangram2-CCC’s performance under varying conditions (e.g., effect strength, spot count, directionality). While synthetic data carries caveats around realism, they enable precise, systematic evaluation.

Using public scRNA-seq data of human breast cancer ^39^, we generated 864 synthetic datasets with artificial interactions, varying in the direction (up or down) of effect gene regulation, baseline gene expression levels, the number of spots, and the strength of up/down-regulation

(**Methods**). Results (see **Figure 3D** and **Supplementary Figure 6**) show that Tangram2 reliably detects effective genes regardless of whether they are up- or downregulated. As expected, the method’s performance improves with more spots (*i*.*e*., more interaction events), and with larger effect sizes (*i*.*e*., stronger signal). In addition, Tangram2 can detect both up- and down-regulated genes, with slightly better performance on the up-regulated ones. The number of effect genes does not influence performance in this analysis. Notably, we only evaluated independent effect genes; however, due to our projection regularization, having more effect genes with shared dependencies could theoretically improve performance. The clearest observation is that it is more difficult to detect changes in genes with low baseline expression or high dropout rates. We speculate this is because there are very few non-zero instances available for scaling when generating the synthetic data. However, detecting signals is difficult when baseline gene expression is very low due to dropout.

### Tangram2 reveals the immunosuppressive mechanisms in the tumor microenvironment of TNBC

After having validated each Tangram2 module, we use Tangram2 to characterize the immunosuppressive paths in the tumor microenvironment of TNBC. TNBC is the most aggressive type of breast cancer and, unlike other types such as the ER+ (estrogen-receptor positive), its microenvironment is characterized by a paradoxically immunogenic tumor microenvironment: it shows high immune cell infiltration, yet these immune responses are actively suppressed ^33,56^. The goal of our analysis is to reveal the mechanisms underlying immunosuppression. The dataset we used for the analysis comprises 10X Visium data from two TNBC patients and two ER+ patients as well as scRNA-seq of more than 75,000 cells from 11 TNBC patients and 10 ER+ patients ^39^. We begin by applying Tangram2-mapping to integrate the scRNA-seq and SRT of both cancer types (**Methods; Supplementary Figures 5)**.^57^ From the integrated data, we compute the average colocalization matrix for each cancer type (**Methods**; **Figure 4A**). Expected cell-cell colocalizations are found for blood vessel units, which include perivascular-like and endothelial cells (PVL Immature and endothelial ACKR1/RGS5 in TNBC;

), as well as for immune aggregates, which are enriched for B-cell and Helper CD4+ T-cells (B-cell Memory/Naive and T-cells CD4+ for both TNBC and ER+). Note that, as TNBC tumors are typically inflamed ^32^, immune cells and CAFs are in proximity of the cancer cells. In contrast, for ER+ breast cancer, we observe that cancer cells form their own aggregates, with negative colocalization relationships with immune cells, indicating they are either surrounding the cancer region or scarcely present. The lower immune infiltration of ER+ cancer as compared to TNBC is consistent with literature^56^.

Next, we apply Tangram2-CCC focusing on identifying the immunosuppressive effects of various cell types in TNBC. **Figure 4B** shows multiple immunosuppressive mechanisms in the tumor microenvironment that affect the CD8 T cell effector functions, described in detail in the following sections and **Figure 4C**.

#### 1 Tumor cells suppress the immune response of CD8 T cells directly through PD-L1 expression

PD-L1 is an immune checkpoint gene constantly found upregulated in the tumor microenvironment, which assists cancer cells to evade immune surveillance ^58,59^. We subcategorised cancer epithelial cells into *PDL1*+ and *PDL1*-populations, based on expression of PD-L1 gene (**Methods**). To investigate the cellular communication effects between CD8^+^ T cells and PD-L1–enriched cancer cells, we compute difference of interaction coefficients, obtained with Tangram2-CCC, between CD8 T cell ← *PD-L1*+ cancer and CD8 T cell ← *PD-L1*-cancer (**Figure 4C(1)**). We observe a clear signature of T cell exhaustion (a dysfunctional state where T-cells lose their effector function), characterized by the upregulation of the master exhaustion regulator *TOX* and several co-inhibitory receptors, including *PDCD1, LAG3, CTLA4, TIM-3* (*HVCR2*), and *TIGIT* ^*58,59*^. Concurrently, there is a significant downregulation of T cell cytotoxicity markers such as *IFNG* and *IL2*. The collective shift corroborates the critical role of PD-L1 in suppressing anti-tumor T cell responses, leading to T cell exhaustion and impaired cytotoxic activity ^60–63^. The binding of *PD-L1* on tumor cells to *PD-1* on CD8 T cells delivers an inhibitory signal that reduces T cell proliferation, cytokine production (e.g., *IFN-γ*), and cytotoxic potential. Interestingly, we observed, upregulation of the marker genes that characterize hypoxia, including *HIF1A, HIF2B* (*ARNT*), *HIF2A* (*EPAS1*), as well as the gene *VEGFA*, suggests that these cells could be in a hypoxic microenvironment, which is known to be prevalent in TNBC^64 65^.

#### 2 Regulatory T cells (T-regs) also plays a critical role in suppressing CD8 T cells

Regulatory T-cells (T-regs) are potent suppressors of CD8 T cell function in the tumor microenvironment. Consistent with the known functions of T-regs, conventional co-inhibitory receptors and T cell exhaustion markers (*TOX, CTLA4, PDCD1, LAG3, HAVCR2* and *TIGIT*) are upregulated in CD8 T cells upon interaction with T-regs^58,59^. The interaction coefficients of hypoxia markers, including *HIF1A, HIF2B* (*ARNT*), *HIF2A* (*EPAS1*), *VEGFA*, however, are insignificant in the CD8^+^ T cell ←T-reg interactions, in contrast to its induction during CD8^+^ T cell ← PDL1+ cancer interactions. Since these marker genes are typically upregulated in response to hypoxia ^66^, this suggests that CD8^+^ T cell ←T-reg interactions are distinct from the hypoxic response where PD-L1^+^ tumor cells interact with CD8^+^ T cells. Given that *HIF1A*, like PD-L1, is known to influence T cell function in the context of chronic antigen stimulation ^67,68^, these interactions may drive subtly different exhaustion phenotypes in CD8^+^ T cells. This finding supports the hypothesis that CD8^+^ T cell ←T-reg and CD8^+^ T cell ← PDL1+ cancer interactions, while both promoting exhaustion, may do so through mechanistically distinct pathways.

#### 3 Cancer cells and macrophages in the tumor microenvironment further boost the immunosuppressive response of T-regs

Building on the previous section, where CD8^+^ T cells engaging T-regs upregulate multiple inhibitory receptors, we next examined which T-reg mechanisms are preferentially engaged in the TNBC tumor microenvironment. T-regs can suppress anti-tumor activities through multiple mechanisms: (i) direct cytotoxicity against effector T cells via *GZMB*, ii) secretion of immunosuppressive cytokines (e.g. *IL10*) (iii) attenuation of antigen-presenting-cell (APC) function via inhibitory check points (*CTLA4, LAG3* and *TIGIT*). From our Tangram2 analysis (**Figure 4C**), T-regs in neighborhood enrichment for tumor cells and macrophages show up-regulation of *CTLA4, LAG3, TIGIT*, down-regulation of *GZMB* and almost unchanged level of *IL10* expression. The pattern is consistent with a shift towards APC-directed suppression (mechanism iii), offering a mechanistic link to the checkpoint-dominant exhaustion we observed in CD8^+^ T cells above. Overall, T-regs adjacent to macrophages and tumor cells display an enhanced, checkpoint-dominant suppressive program, in line with prior reports of high checkpoint expression on T-regs in the tumor microenvironment. ^69,70^

#### 4 The TME seems to reprogram macrophages towards an immunosuppressive phenotype

Based on Tangram2 analysis (**Figure 4C**), macrophages adjacent to cancer epithelial display downregulation of the *MHC II* genes such as *HLA-DRA, HLA-DRB1* as well as *CCR7*, indicating reduced antigen representation and impaired lymph-node-directed migration. ^71–73^ In parallel, the B7 landscape shifts toward increased *CD80* and decreased *CD86*. Given the elevated *CTLA4* on T-regs and CD8+ T cells (from previous sections), this *CD80*^high^/*CD86*^low^ configuration is poised for *CTLA4*–mediated removal of B7 ligands, thereby attenuating *CD28* co-stimulation of effector T cells and reinforcing checkpoint-dominant suppression ^74,75^. Collectively, the analysis supports a model in which cancer–macrophage crosstalk drives tumor-associated macrophages toward more immunosuppressive states.

#### 5 CAF-mediated immunosuppression in the TNBC microenvironment: a potentially complex interplay

Cancer-associated fibroblast (CAFs) represent a critical component of the tumor microenvironment (TME), actively contributing to tumor progression, metastasis and therapeutic resistance. CAFs possess multi-faceted roles in driving tumor growth and immunosuppression through diverse mechanisms, including the secretion of growth factors, extracellular matrix (ECM) remodeling, and metabolic alterations ^76,77^. In this section, we investigated the interaction between CAFs and cancer epithelial cells in the two TNBC patients. **Figure 4C(5)** shows the interaction coefficients of growth factors and immunosuppressive genes enriched in cancer epithelial cells upon interactions with CAFs. Among the growth factor genes, the significant upregulation of *TGFb1* and *HGF*, alongside with the modest upregulation of *PDGFB, PDGFC* and *IGF1R*, indicate mechanisms by which cancer cells drive proliferation and invasion through cellular crosstalk with CAFs^78–81^. Furthermore, although the interaction coefficient of *CD274* (PD-L1) is close to 0, the increased expression of other immune checkpoint and regulatory molecules, including *PDCD1LG2* (*PD-L2*), *CD276 (B7-H3), TNFRSF14, C10orF54* (*VISTA)* and *LILRB2*, indicate that immunosuppressive tumor cell states can be maintained or promoted by CAFs ^82^. PD-L2, for example, processes high affinity to PD-1, enables T cell inhibition and contributes to immune escape, while B7-H3 may also hold similar immunoregulatory properties^83,84^. The observed profiles point to cancer cell growth and immune-evasive tumor phenotypes induced by cancer-CAF interactions, which provide additional mechanisms to the direct and indirect paths of tumor immune suppression identified in Figure 4D. This finding underscores the critical and intricate role of CAFs in shaping the tumor microenvironment, which indicates a need for further investigation and potentially multi-target or combination approaches in cancer therapeutics.

In conclusion, a number of immunosuppressive mechanisms within the tumor microenvironment of TNBC have been recapitulated. We show that tumor cells suppress the effector function of CD8 T cells either directly through expressing ligands such as PD-L1, or indirectly through interaction with other cell types such as regulatory T cells and macrophages. Furthermore, CAFs support cancer cell growth and additionally shape cancer immunosuppressive states. Tangram2 enables better understanding of cellular interactions in the tumor microenvironment, which potentially helps to better identify therapeutic targets.

### Tangram2-CCC uncovers survival-linked interactions in the cSCC tumor microenvironment

Having validated Tangram2 across diverse settings, we next applied it to a larger dataset to uncover new insights into cell-cell interactions. We selected a publicly available cutaneous squamous cell carcinoma (cSCC) dataset from Ji et al (dataset ID: cscc), which consists of paired scRNA-seq and Visium data from 6 patients, with 2-3 Visium replicates per patient. cSCC progression is known to be driven by complex tumor microenvironment interactions, particularly among immune cells such as macrophages and regulatory T cells ^34,85^. These interactions foster an immunosuppressive environment that supports tumor growth and resistance. Characterizing these dynamics may uncover new opportunities for therapeutic intervention.

We narrowed down our analysis to the Macrophage←Treg interaction, and identified the genes most upregulated upon this interactions (**Methods)**. The Macrophage←Treg gene set included 259 genes, with *STAT1, ISG15, ISG20, IFITM3, CXCL9* and *CXCL10*—key in immune cell recruitment and anti-tumor activity—ranking in the top 10, enriched for interferon responses. Macrophages expressing interferon-stimulated genes (*e*.*g*., *ISG15, ISG20*) and T cell–recruiting chemokines (*e*.*g*., *CXCL9, CXCL10*) exhibit a canonical type I interferon-driven pro-inflammatory phenotype. Hence, our results seem to suggest that macrophages interacting with Tregs exhibit an M1-like, pro-inflammatory phenotype with elevated cytokine levels. This finding contrasts with prior reports that Treg–macrophage crosstalk typically promotes an M2-like, immunosuppressive phenotype ^86,87^. One plausible explanation is that the tumor-infiltrating Tregs in our cSCC patients express high levels of GITR (**Supplementary Figure 10**), which can reprogram them toward a Th1-like effector state and induce interferon (IFNγ) production, as shown in recent studies on *GITR* signaling in Tregs. ^88–91^ The resulting IFNγ-rich microenvironment could, in turn, drive macrophage polarization toward an M1-like phenotype. The M1 macrophage state is often associated with more positive outcomes for cancer patients, and has in some cases even been implicated to be necessary for the efficacy of the PD-L1/PD-1 blockade, meaning this Macrophage ← Treg interaction could be highly desirable in the cSCC cancer ^92–95^.

To explore the potential therapeutic relevance of this interaction, we analyzed it in patients with Skin Cutaneous Melanoma (SKCM) using data from TCGA—the most suitable available substitute for cSCC within the TCGA dataset. The Spearman correlation between *GITR* expression and macrophage←Treg enrichment was strong in SKCM (r = 0.651, p = 8.8 × 10^-53^), see **Supplementary Figure 11**. This pattern was not observed in Lung or Head and Neck Squamous Cell Carcinomas, suggesting the signal may be specific to skin cancers. Patients in the highest quartile of macrophage←Treg enrichment score exhibited significantly better survival compared to those in the lowest quartile. Results are shown in **Supplementary Figure 12**.

While the strong correlation between macrophage←Treg enrichment and overall immune infiltration is noted, it remains unclear whether the observed phenomenon has a causal effect on the survival response or is merely an association to the pre-existing anti-tumor response. Nevertheless, these findings propose the macrophage-Treg interactions as a potential clinical target, which merits further investigation. Though additional validation is beyond this study’s scope, this work exemplifies Tangram2’s utility in target discovery using public datasets.

### Tangram2 reveals the immunosuppressive activities of tumor-specific keratinocyte (TSK) in cutaneous squamous cell carcinoma (cSCC)

Besides the interactions between immune cells, the tumor-specific keratinocyte (TSK) population is a unique subpopulation of tumor cells in cutaneous squamous cell carcinoma (cSCC). Based on earlier human clinical trials of anti-PD-1 treatments, high expression of TSK-specific genes such as *ITGB1* and *PLAU* is associated with reduced progression-free survival, potentially due to either the immunosuppressive activity of TSKs or its intrinsic resistance to immune attack ^50,96^. Tangram2 enables investigating the potential immunosuppressive activities of TSK using Tangram2 (**Figure 5A**).

**Figure 5.**
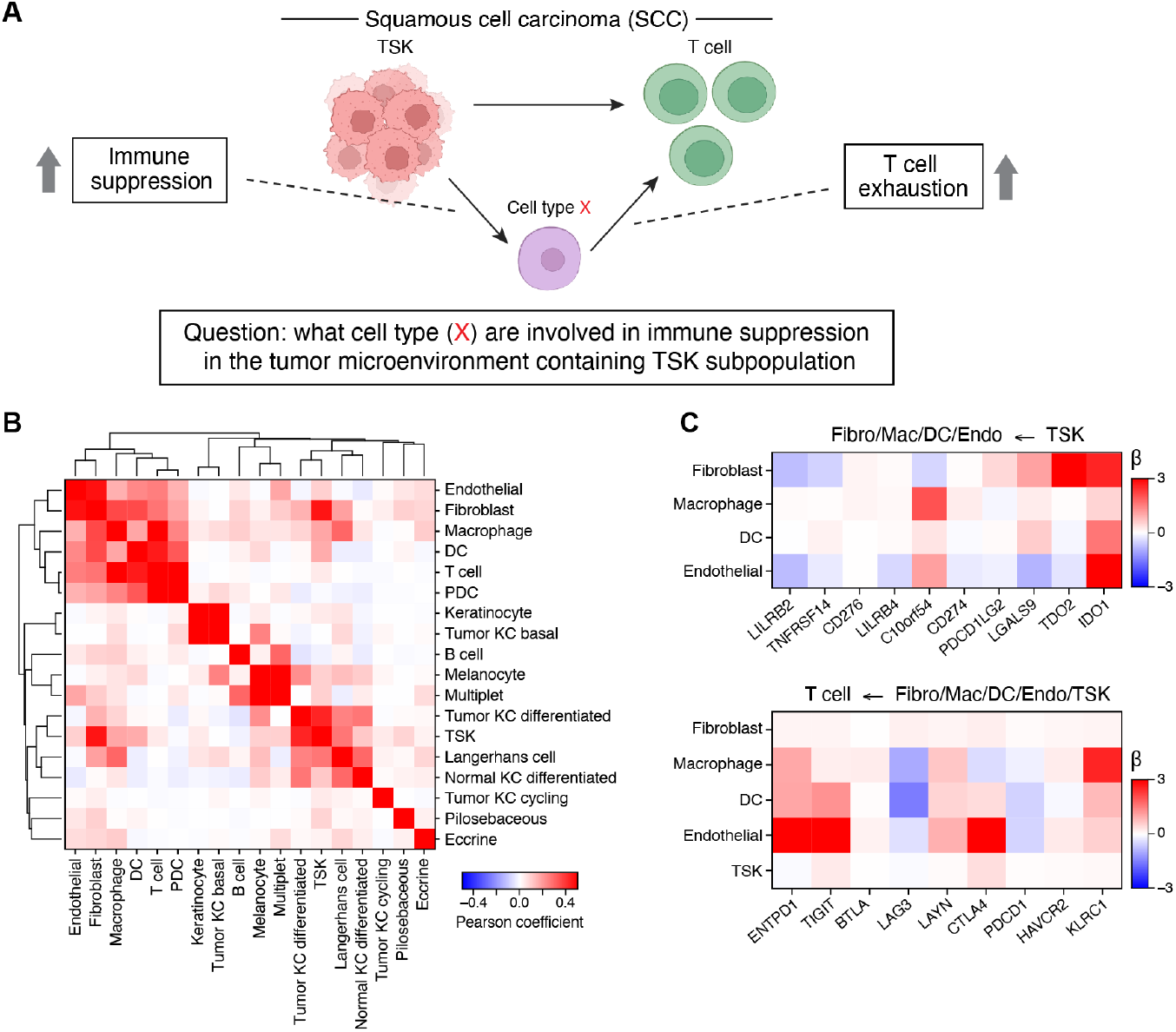
Tangram2 reveals cellular interaction relationships in cutaneous squamous cell carcinoma (cSCC), offering novel biological insights. **A**. Scheme diagram of the cSCC data study: investigating the immunosuppressive paths in the tumor microenvironment containing TSK and the T cells; **B**. Cluster maps of the average colocalization scores between cell types of cSCC samples based on the Tangram 2 mapping. The cell types that consist of less than 0.05% of total cell populations in the spatial samples are excluded from this analysis **C**. The interaction coefficients of selected gene list learned by Tangram2-ccc of the interaction *fibroblast/ endothelial/ macrophages/ DC <– TSK* and *T cells <– fibroblast/ endothelial/ macrophages/ DC/ TSK*. The selected gene lists are either immunosuppressive markers or T cell exhaustion markers given in the cSCC paper ^50^.

Compared to other tumor subtypes (such as differentiated, basal-like and cycling tumor cells), TSK shows significantly stronger colocalization patterns with endothelial cells, fibroblast, DC, macrophages and T cells, while these cell types also strongly colocalize with T cells according to Tangram2-mapping analysis (**Figure 5B)**. The colocalization pattern suggests the possibility of both a direct immunosuppressive path between TSK and T cells, and indirect paths mediated through these four different cell types.

Similar to the TNBC analysis in **Figure 4B**, we investigate various immunosuppressive mechanisms between TSK and T cells through Tangram2-CCC. **Figure 5C** presents the median interaction coefficients of immunoinhibitory markers on T cells, calculated across all five patients, upon interaction with TSK, endothelial cells, fibroblasts, dendritic cells (DCs) and macrophages. Consistent with analysis in the original paper in squamous cell carcinoma (SCC)^50^, we focused on immunosuppression-associated genes for interactions where DCs, fibroblasts, macrophages, and endothelial cells were receiver cell types, and co-inhibitory and exhaustion markers when T cells were the receiver in **Figure 5C**. For the T cell ← TSK interaction, the interaction coefficients of T cell exhaustion and co-inhibitory markers, including *PDCD1, LAG3, TIGIT*, are close to 0 (the last row of lower panel of **Figure 5C**), indicating that there are no strong direct immunosuppressive effects from TSK cells on T cells. In contrast, significant immunosuppressive activities are observed from the indirect paths (**Figure 5C**). The elevated expression of metabolic enzymes (*IDO1, TDO2*) in the stromal cells upon interaction with TSK, can result in suppression of T cell proliferation and promotes T-regs differentiation ^97,98^. Furthermore, the upregulation of immune checkpoint molecules in stromal cells including VISTA (*C10orF54*), *LGALS9* and PD-L2 (*PDCD1LG2*), and the upregulation of inhibitory markers in T cells such as *TIGIT, CTLA4* and *KLRC1*, further compromise the anti-tumor response in the tumor microenvironment ^59^. These findings confirm the immunosuppressive activity of TSKs, which potentially lead to its association with reduced progression-free survival in a clinical trial of *PD-1* checkpoint inhibitor treatment.

## DISCUSSION

Methods integrating single-cell RNA sequencing (scRNA-seq) and spatial transcriptomics (ST) have become indispensable for mapping cellular organization within tissues, underpinning large-scale efforts such as the Human BioMolecular Atlas Program (HuBMAP) ^99^ and the Human Cell Atlas (HCA) ^100^. Here, we refine this paradigm with Tangram 2 and extend it to learn cell–cell communications (CCCs) directly from data.

A complete understanding of cellular interactions requires not only transcriptomic profiles but also the spatial and environmental context in which cells reside. This has motivated the growing integration of scRNA-seq with ST data. However, current ST platforms still face trade-offs between spatial resolution and transcriptome coverage, making integrative computational frameworks essential for capturing communication mechanisms at scale. Emerging high-resolution, high-throughput ST technologies may mitigate these limitations, but their accuracy and reproducibility remain variable at present. However, as these technologies mature, they will likely reshape the landscape of computational CCC inference.

A persistent challenge in CCC analysis is the lack of systematic benchmarking. Many tools assess performance by reproducing known biology, which limits objective comparisons and hinders method selection. In Tangram 2, we implemented rigorous benchmarking across diverse datasets and evaluation schemes before deriving biological insights, providing a foundation for reproducible and transparent assessment.

By revealing intercellular communication programs that shape tissue organization and function, Tangram 2 offers a robust framework for studying cell–cell interactions. Its application to triple-negative breast cancer and squamous cell carcinoma illustrates its potential to uncover therapeutic targets and advance mechanistic understanding of complex cellular ecosystems. Looking ahead, extending Tangram 2 to integrate additional modalities and species will further enhance its ability to model communication across biological systems.

## METHODS

### Tangram2-mapping Description

The matrix 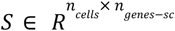 denotes the cell by gene expression from scRNA-seq, where n_cells_ is the number of cells in the scRNA-seq and n_genes-sc_ is the number of genes measured in the scRNA-seq. *S*_*ik*_ provides the expression level of gene k of cell i. The matrix 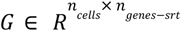 symbolizes the spot-by-gene expression from spatial transcriptomics, where G_j*k*_ denotes the expression level of gene k at spot j. There is also a 1D vector 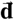. The dimension of 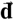 is equivalent to the number of spots measured in ST data while the value indicates the density of transcriptome of each spot where ∑*d*_*j*_=1.

Building on the framework of its predecessor, Tangram2-mapping is designed to construct a spatial alignment (indicated by matrix T) between single-cell and spatial data. Here, T_ij_ denotes the probability of cell i in location j. Tangram2-mapping offers two distinct mapping modes, each tailored to accommodate different data conditions and discussed in the succeeding section. The optimization of matrix T is dependent on the selected mapping mode, thereby ensuring the most precise alignment specific to the given respective data scenario.

### Tangram2-mapping Modes

#### Tangram2-mapping cell-level mode

This mode is employed when single-cell and spatial transcriptomics data originate from the paired dataset, characterized by similar cell type compositions. In order to align the scRNA-seq 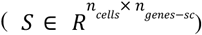 with the spatial transcriptomics 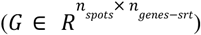, a mapping matrix 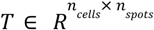 is learned, as driven by the loss function. The loss function of the cell-level probabilistic mapping mode is given by:

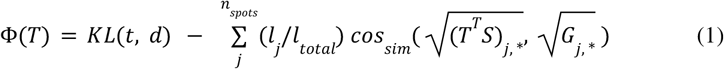

Here *KL* indicates the Kullback-Leibler divergence, where the 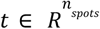 and 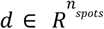 are predicted and measured cell density that satisfies 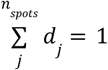 and 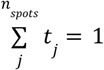. The vector *t* is calculated from the mapping matrix *T* by 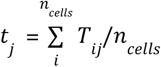. The cos_sim_ signifies the cosine similarity function, calculated by 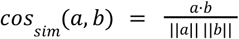 and * is the matrix slicing. The *l*_j_ in Equation (1) refers to the number of transcriptomes measured in spot j, while the *l*_total_ refers to the total number of transcriptomes measured in ST sample. In Tangram2-mapping, a soft assignment T is achieved between the scRNA-seq data and spatial transcriptomics data. The T_ij_ refers to the probability of the cell i mapped to spot j. Given this definition, there is a probability matrix constraint T such that 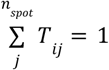 and *T*_*ij*_ > 0. These constraints are enforced by introducing a softmax function along the spot axis for any given 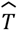, such that:

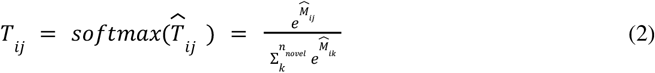

By minimizing the objective function with the constraints presented in equation (2) using Adam optimizer, Tangram 2 intuitively adapts an optimal alignment matrix T. From this alignment matrix T, one can derive the probabilistic distribution of diverse cells/cell types, while the T^T^S provides the predicted gene distribution.

### Tangram2-mapping cluster-level mode

The cluster-level probabilistic mapping mode is primarily designed for unpaired data, enabling the utilization of unpaired open-source scRNA-seq datasets. Rather than finding the probability assignment of each individual cell to a spatial location, the model initially computes the mean expression for each cell type. The matrix 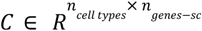 is computed from S by calculating the average gene expression among cell types, where n_cell-type_ is the number of cell types in the scRNA-seq. The alignment matrix 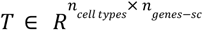 from the cluster-level mapping characterizes the spatial distributions of cell type. Furthermore, there is an additional vector **w** (dimension: the number of cell types), where w_i_ indicates the proportion of cell type i in the spatial sample. Subsequently, the mapping matrix T and the corresponding cell type ratios 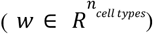 are jointly inferred to maximize the alignment between the mapping result (wT^T^C) and the spatial gene expression data (G). The objective function of cluster-level probabilistic mapping mode is as followed:

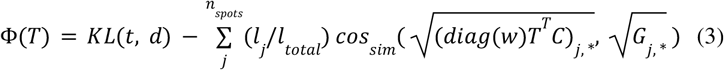

Here the diag(w) refers to the diagonal matrix formed by vector ***w***. *l*_*j*_ is the number of transcriptomes at spot j, while *l* _*total*_ refers to the total number of transcriptomes. In addition to the constraint on the probabilistic alignment matrix ***T***, there is a constraint on the cell type composition vector w, such that 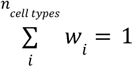 and *w*_*i*_ > 0. The constraint is also enforced by the softmax function:

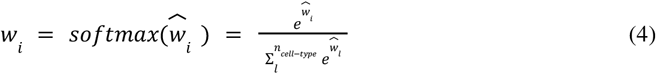

By optimizing the objective function subject to the dual constraints detailed in Equations (2) and (4), Tangram2-mapping efficiently yields the optimal alignment matrix, T, in conjunction with the cell type compositions, denoted by ***w***. The distribution of each cell type can be calculated from (*diag*(*w*)*T*^*T*^).

### Tangram2-mapping integrate mode

As its name indicates, the integrated mapping combines the cluster-level and cell-level mapping, such that it allows mapping of unpaired datasets while preserving the cell-level information. The integrated mapping first applies the cluster-level mapping, from which the cell type composition ***w*** is learned. After that, the scRNA-seq data is normalized using ***w***. The renormalization step takes the following form:

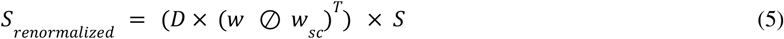

Here ***w***_sc_ is the proportion of each cell type in the single cell datasets calculated based on cell ount. 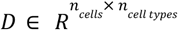 is a cell type indicator matrix where where *D*_*ik*_ = 1 if cell *i* belongs to cell type *k* and zero otherwise, a cell can only belong to one type, i.e.,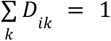. The ⊘ refers to the element-wise division between two vectors of the same length.

After normalization, the single cell (*S*_*renormalized*_) and spatial transcriptomics (G) are expected to have similar cell type compositions. Then cell-level mapping is applied to compute the alignment matrix T, which is the probability assignment of each normalized cell to spatial locations.

### Synthetic Data Generation

To generate synthetic data we employed a strategy similar to what has been presented in multiple works; where cells from single cell data are distributed to artificial spots. However, our approach distinguishes itself in several key ways: i) it supports spatial coherence of cell type populations, ii) it supports the introduction of artificial cell-cell interactions and downstream effects, iii) it supports the introduction of multinomial noise. Below we describe the base generation approach and then elaborate on how these features are incorporated into our synthetic data.

### Data Generation

Let *S*_*seed*_ represent the “seed” single-cell dataset from which we aim to create a new tuple (*S, G*) of synthetic data. The user specifies the following parameters: *n_spots* (number of spots), *n_cells_per_spot* (expected number of cells per spot), and *n_types_per_spot* (expected number of cell types per spot). Based on these inputs, we generate *n_spots* artificial spots. For each spot, we sample the number of cells and cell types from a Poisson distribution, using *n_cells_per_spot* and *n_types_per_spot* as the means. The number of cell types per spot is limited to a minimum of 1 and a maximum of the total number of available cell types.

We then randomly assign cells from *S*_seed_ to these artificial spots, and the gene expression of each spot is defined as the sum of the gene expression of all cells assigned to that spot. The resulting synthetic data tuple (*S, G*) consists of the generated spatial data (*G*) and all cells (*S*) used to construct the spatial data. This pairing is exact, as the assignment of cells to spots is fully described by the matrix *T*, with *G* = *T*^*T*^S. We also retain the matrix *N*, which indicates the number of cells from each cell type in every spot.

### Noise

We add what we call “multinomial noise” to the data, this noise operates on the raw (count) spatial data. For each spot *s* we compute the proportion vector p_s_ where p_sg_ indicates the proportion of transcripts that belong to gene *g* in spot *s*. Let *t*_*s*_ be the total number of transcripts in spot *s*, and *m*_s_ a multiplier (default: 1) that indicates what proportion of *t*_*s*_ that should be resampled. Let *ε* be the noise level (default: 0) and 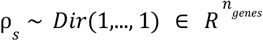 is the noise vector. We then generate noisy data accordingly 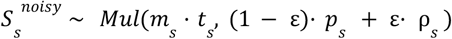

### Spatial coherence

In certain tissues, gene expression demonstrates spatial coherence, a key assumption in many spatial mapping methods (e.g., SpaOTsc). We account for this in our synthetic data generation through the following steps. First, from the matrix N, which represents cell types per spot, we compute a proportion matrix P, indicating the fraction of each cell type at each spot. We then apply principal component analysis (PCA) to project P into a 2D space, i.e., 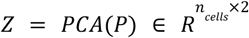. Finally, we use the Earth’s Mover Distance (EMD) to calculate a transport plan M, which relates each cell type proportion profile to the spatial coordinates. We rearrange the spots according to the transport plan. This ensures that spots close to each other maintain similar cell type proportions, preserving spatial coherence. See **Supplementary Figure 7** for a visualization of the synthetic data with and without spatial encoding.

### Synthetic interactions and effects

Each interaction comprises one signal gene and a set of associated effect genes. Cells are divided into three classes: signalers, receptive cells, and non-receptive cells. These classes are mutually exclusive. Signaler cells overexpress the signal gene and activate nearby receptive cells, while non-receptive cells remain unaffected. All cells can express both the signal and effect genes, but expression levels vary depending on the cell class. The assignment of signalers and receptive cells can be restricted to specific cell types or applied to all cells.

To designate signalers, spots are randomly assigned as active or inactive to maintain spatial consistency, with the fraction of active spots determined by a preset probability (default: 0.5). Receptive cells are activated based on a probability (p_inter) and must be located in spots where signalers are present. Finally, synthetic gene expression is introduced for both signal and effect genes.

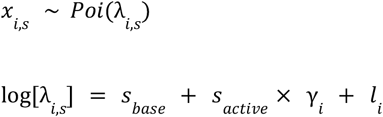

and

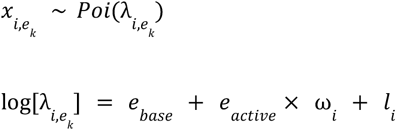

with

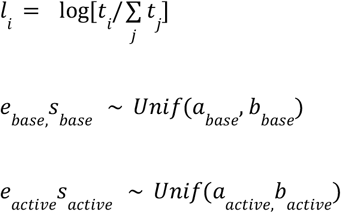

Where *x*_*i,s*_ is the expression of signal *s* in cell *i*, and 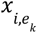 is the expression of effect *k* in cell *i*. Here the coefficients with the “base” subscript are the baseline expression of the signals and effects, the “active” coefficients regulate the impact of a cell being a signaler or receiver on the gene expression. The value of γ_i_ is 1 if cell *i* is a signaler cell and 0 otherwise, ω_i_ is 1 if a cell is an active cell or not. Again, *t*_*i*_ is the total number of transcripts in cell *i*. See **Supplementary Figure 8** shows an example of the distribution of active receivers and signalers as well as the signal and effect gene expression. The cells overall identity is not impacted by the addition of effect genes, see **Supplementary Figure 9**.

### Deconvolution Evaluation

The evaluation of Tangram2-mapping deconvolution is conducted using simulated paired and unpaired datasets (depending on whether the scRNA-seq and SRT are from the same patient) generated by Tangram2-evalkit. These datasets represent three distinct scenarios, reflecting different experimental conditions. The schematic diagram about how the datasets are generated can be found in **Supplementary Figure 1**. Three possible scenarios are shown as follows:

1. *<u>Perfectly Paired dataset</u>*: the spatial dataset is directly generated from the reference single cell data, meaning that each cell in the single cell data can be found at a certain spatial spot. It is to be noted that this is an ideal but not realistic setting.
2. *<u>Paired datasets</u>*: Both the spatial and scRNA-seq datasets are derived from the same tissue of the same patient. While the specific cells in each dataset aren’t identical, they share similar cell type compositions. This scenario reflects a more realistic experimental setting where both datasets are available from the same source.
3. *<u>Unpaired datasets</u>*: The spatial dataset is from one patient, while the scRNA-seq data is from a different patient. This scenario is the most challenging, as it’s possible for the cell type compositions of the two datasets to differ. This represents a common real-world situation where single-cell and spatial data come from different sources.

We focused our evaluation on the paired and unpaired datasets (Scenario 2 & 3), as these scenarios best reflect actual experimental conditions.

The methodologies to generate these datasets of different scenarios from the framework are elaborated in **Supplementary Figure 1B**. The scRNA-seq dataset used here is the triple negative breast cancer (TNBC) data published by Sunny et al ^39^. We selected scRNA-seq data from a single patient (Patient A) and randomly divided it into two halves, A_1 and A_sc_2. We then used Tangram2-evalkit on A_1 to generate both scRNA-seq data (A_sc_1) and a corresponding spatial dataset (A_sp).

1. For the **paired scenario**, we used A_sc_2 as the reference for deconvolving A_sp. In this case, the datasets are from the same patient and have comparable cell type compositions but do not contain the exact same cells.
2. For the **unpaired scenario**, we used scRNA-seq data from another TNBC patient (**X_sc**) as the reference for deconvolving A_sp.

Across these scenarios, the simulated datasets were constructed so that each spot contains up to 10 cells, with a maximum of three different cell types per spot.

### Additional Deconvolution Analysis

In addition to the deconvolution analysis on simulated datasets, we also benchmarked Tangram2 against other deconvolution tools using published datasets. The datasets, benchmarking methodology, along with the accuracy score computation, have been developed by Li et al in their benchmarking paper ^27^. The benchmarking model generates a final accuracy score based on the ranking of four evaluation metrics: Pearson Correlation Coefficient (PCC), Cosine Similarity (CS), Root-Mean-Square Error (RMSE), and Jensen-Shannon Divergence (JS). The detailed information of the benchmarking method and dataset can be found in Li et al’s paper ^27^.

### Evaluation of Mapping

Similar to deconvolution analysis, the evaluation of mapping is also performed using simulated datasets generated by the Telegraph framework. The scRNA-seq datasets for mapping evaluation are derived from the cSCC patients ^50^. Since the ground truth mapping locations of each cell from scRNA-seq are required, the mapping accuracy is assessed using scenarios of *<u>perfectly paired datasets</u>*, as illustrated in **Supplementary Figure 1B**. To introduce mismatches between the SRT datasets and reference scRNA-seq, noise parameters ranging from 0.0 to 1.0 are applied. In addition to the publicly available mapping methods, two baseline approaches are also included for benchmarking:

1. *<u>The random baseline</u>*: assigning each cell to a random spot;
2. *<u>The argmax baseline</u>*: assigning cells to the spot with highest Pearson correlation of gene expression.

An additional parameter to vary is the total number of SRT spots created by Tangram2-evalkit, which ranges from 100 to 500 spots. Mapping becomes more challenging as the number of spots increases. Since most of these methods output a cell by spot probabilistic mapping matrix, the performance is evaluated by Jaccard score obtained from assigning each cell to the spot with highest probability.

### Tangram2-CCC Method

Tangram2-CCC is our cell-cell communication method, in essence it is a way to leverage the cell-to-location maps (T) that are the product from any type of mapping method to infer cell-cell communication effects. We will first clarify a few terms and then describe the method.

### Cell-cell communication and effects

Let A and B denote two cell types (not necessarily distinct), we use the notation A ←B to denote the interaction event where cells of cell type A acts as a receiver and cell type B as a sender of some signal. In the context of a ligand-receptor interaction, A would express the receptor and B the ligand. We define *cell-cell communication effects* as the downstream genes in the receiver cell that are associated with a particular cell type interaction event.

### Model

For a given dataset, assume that we’ve obtained a map 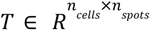 that relates cells to spots, retrieved using some mapping method (e.g., Tangram2). Let 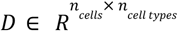 be a cell type indicator matrix, where *D*_*ik*_ = 1 if cell *i* belongs to cell type *k* and zero otherwise, a cell can only belong to one type, i.e., 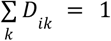.

In the first step we compute the cell-cell adjacency matrix 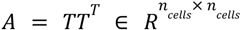, this represents the colocalization of cells based on the mapping. More specifically, *A*_*ij*_ indicates the level of colocalization that cell *i* and cell *j* exhibit. Next, we compute the effective neighborhood matrix 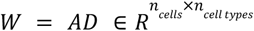. This matrix presents the effective cell type neighborhood of each cell based on the mapping. That is, *W*_*ik*_ indicates the abundance of cell type *k* in the neighborhood of cell *i*.

In our second step, we will use *W, D* and a normalized version of the single cell gene expression matrix (*S*). We normalize *S* using a standard procedure, for each cell we: i) divide all counts by the total sum of counts in that cell and multiply all values with 10^4^, ii) apply a log1p transformation to the result. We let *X* denote the normalized single cell gene expression. We consider the gene expression of a gene *g* in a particular cell *i* to be mainly dependent on three factors: i) the inherent baseline expression of that gene (shared across all cells), ii) the cell type of the cell, iii) the environmental context of the cell (represented by the effective neighborhood) – how the environment impacts the gene expression will depend on the cell type of the cell. This results in the following model

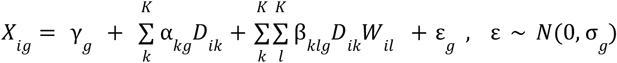

Which can be written as:

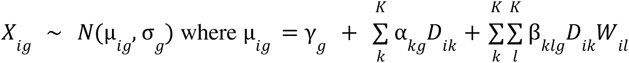

In this instance (γ, α, β, σ) are all learnable parameters, *K* is the number of cell types. We will learn the value of these parameters using maximum likelihood estimation (MLE). Our loss function, for each gene *g* will be 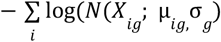 that is, the negative log-likelihood of observing *X*_*ig*_ given the value of μ and σ. As is obvious from the model, this requires one model per gene, resulting in 2*G* + (*K* × *N*) + (*N* × *K* × *K*) parameters and a computational complexity of *O*(*NK* ^2^) – *N* being the number of genes and *K* the number of cell types. In transcriptomics data then number of genes is often in the magnitude 10^4^ while the number of cell types are, usually, in the range of 1-50 meaning that despite *K* being quadratic, the dominant factor is the number of genes N. To reduce computational time we thus implement a computational trick, where we fit our model in a lower dimensional space and then project the parameter values back to the original space.

We can let *X* ≈ *Q* where 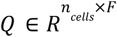 and *V* ∈ *R*^*F*×*G*^, here V are the *F* first PCA components and *Q* is the associated PCA loadings. We replace *X* in the previous model with *Q*, and fit it using the same procedure as described; this reduces the complexity to *O*(*FK* ^2^) where F is a number chosen by the user, our default is 500. This results in a set of learn parameters (*Vγ* ^*Q*^, *Vα*^*Q*^, *V*β^*Q*^, *V*σ^*Q*^). Finally, we use *V* to project the learnt parameters to the original domain, that is: (γ, α, β, σ) ≈ (*Vγ* ^*Q*^, *Vα* ^*Q*^, *V*β^*Q*^, *V*σ ^*Q*^). To be noted, some accuracy will be lost since we use an approximation, however we use this for all our analysis and still achieve satisfactory results.

For a given dataset, we assume a map 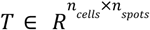 that relates cells to spots, obtained via a mapping method (e.g., Tangram2). Let 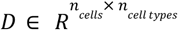 types be a cell type indicator matrix where where *D*_*ik*_ = 1 if cell *i* belongs to cell type *k* and zero otherwise, a cell can only belong to one type, i.e., 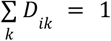.

First, we compute the cell-cell adjacency matrix 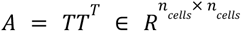, representing the colocalization of cells based on the mapping, where *A*_*ij*_ indicates the colocalization between cell *i* and cell *j*. Next, we compute the effective neighborhood matrix 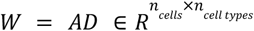 where *W*_*ik*_ indicates the abundance of cell type *k* in the neighborhood of cell *i*. These steps are depicted in Figure 1a and b.

*S* is normalized by dividing each cell’s counts by the total counts, multiplying by 10 ^4^, and applying a log 1*p* transformation. Let *X* denote the normalized expression. We model gene expression *X*_*ig*_ in cell *i* for gene *g* as depending on: i) the gene identity, ii) the cell type, and iii) the environmental context, with the model:

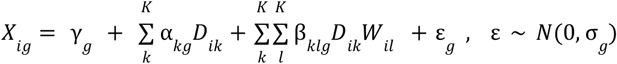

Or equivalently,

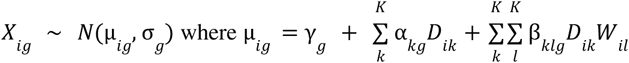

Here, (γ, α, β, σ) are all learnable parameters while *K* is the number of cell types. Briefly, γ _*g*_ is the baseline gene expression of gene *g, α* _*kg*_ is the effect of cell type *k* on gene *g*, and β_*klg*_ is the effect on gene *g* in cell type *k* when cell type *l* interact with it (interaction: k ←l). In the main text, we refer to β as *interaction coefficients*.

We estimate these parameters via maximum likelihood estimation (MLE), with the loss function for each gene *g* being the negative log-likelihood of the normal distribution 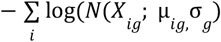. This results in a total of 2*G* + (*K* × *G*) + (*G* × *K* × *K*) parameters to be learnt and a computational complexity of *O*(*GK* ^2^), where *G* is the number of genes and *K* the number of cell types. This is illustrated in Figure 1c.

Since *G* is often on the order of 10 ^4^, while *K* is typically between 1–50, the number of genes dominates the complexity. To reduce computational time, we fit the model in a lower-dimensional space and project the parameters back. Let *X* ≈ *QV*, where 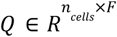 and 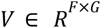 are the first *F* PCA components and loadings. By replacing *X* with *Q*, we reduce the complexity to *O*(*FK* ^2^), where *F* is user-defined (default: *F* = 500). This results in a set of learnt parameters (*Vγ*^*Q*^, *Vα*^*Q*^, *V*β ^*Q*^, *V*σ^*Q*^). Finally, we use *V* to project the learnt parameters to the original domain, that is: (γ, α, β, σ) ≈ (*Vγ*^*Q*^, *Vα*^*Q*^, *V*β^*Q*^, *V*σ ^*Q*^). To be noted, some accuracy will be lost since we use an approximation, however we use this for all our analysis and still achieve satisfactory results.

For the parameter fitting we use Adams as an optimizer, with 1000 epochs and 0.01 learning rate unless otherwise stated.

### Tangram2-CCC Analysis

#### Slide-tags analysis

In the Slide-tags analysis, we “visified” the original data by placing a 25×l25 regular grid over the spatial domain, creating 625 grid points that act as spots or spots. Cells within 50 units of a grid point (in original coordinates) were assigned to a spot, and cells not assigned to any spot were removed. The sum of transcripts in each spot was treated as the spatial gene expression profile. The scRNA-seq data was taken as the same cells assigned to spots but without spatial coordinates.

For pre-processing, we handled the semi-synthetic visified spatial and scRNA-seq data similarly to standard Tangram2-CCC analyses. We removed features starting with “MT-”, “MT”, “RP”, “RPS”, and “LINC” as well as those ending in “.1”. In scRNA-seq data, cells with fewer than 300 total counts and genes with fewer than 10 counts across all cells were removed. For spatial data, cells with fewer than 100 total counts and genes with fewer than 10 total counts were excluded. Highly variable genes were extracted from the scRNA-seq data using Scanpy with the following steps:

- sc.pp.normalize_total(…, 1e4)
- sc.pp.log1p(…)
- sc.pp.highly_variable_genes(…, n_top_genes=5000)

After preprocessing, we ran Tangram2 to obtain the cell-to-location map *T*. Tangram2 was run on raw counts using the highly variable genes as training genes. We used this map, along with normalized gene expression (*X*) and the cell type indicator matrix (*D*), as input to Tangram.

To validate the interaction coefficients, we tested whether the top 10 genes (ranked by the interaction coefficient) were up-regulated in receiver cells when near signaler cells. For each interaction A←B, we calculated the distance between receiver cells (A) and their nearest signaler cells (B) using the original spatial coordinates. We defined proximal and distal receiver cells based on the 25th and 75th quantiles of the distance distribution. For each of the top 10 genes in all cell-type combinations (excluding self-interactions), we computed the difference in mean expression between proximal and distal cells and counted how often the proximal expression was higher.

To assess significance, we performed a binomial test. Let *N* = 1560 be the total number of comparisons (13 cell types × 12 interactions × 10 genes), and *k* the number of successes (higher expression in proximal cells). Under the null hypothesis *p* = 0. 5 (equal likelihood of success or failure), we computed the p-value as Thus, *p*_*value*_ = *P*(*X* ≥ *k*; *n* = 1560, *p* = 0. 5).

#### Co-culture analysis

In the co-culture analysis, we processed the cell culture dataset (dataset ID: cell cultures) to obtain ground truth gene expression profiles. The dataset comprised 94 deposited experiments in which splenic CD11c^+^ dendritic cells (DCs) from C57BL/6 wild-type mice were either cultured alone or co-cultured with OT-II CD4^+^ T cells after antigen loading with OVA and activation with the Toll-like receptor 4 agonist lipopolysaccharide (LPS). DCs were split into two groups based on whether they were grown in mono-culture or co-culture.. Gene expression was normalized using the following steps in Scanpy:

- sc.pp.normalize_total(…, target_sum=1e4)
- sc.pp.log1p(…)

We calculated logfoldchanges and adjusted p-values using the Wilcoxon method in Scanpy:

- sc.tl.rank_genes_groups(…, method=‘wilcoxon’)

Genes with an adjusted p-value ≤ 0.05 and positive log fold change were kept as the ground truth set, see Figure 3C.

For scRNA-seq and Visium data, we used mouse colon data (dataset ID: colon) consisting of four Visium sections and one scRNA-seq dataset. For the Visium data, we concatenated the sections named 1A_Hh, 2A_Hh, 3A_Hh, and 4A_Hh. We applied similar filtering as in the Slide-tags analysis, adjusting gene removals for mouse data by excluding features starting with “mt-”, “Mt”, “ENSMUSG”, “rps”, “rpl”, “rp-”, “rp”, and “Rik”. We harmonized label granularity between the cell culture experiments and colon scRNA-seq by relabeling ‘naive thymus-derived CD4-positive, alpha-beta T cell’, ‘regulatory T cell’, ‘CD4-positive, alpha-beta memory T cell’, and ‘T-helper 17 cell’ as CD4+ cells. Cells labeled as ‘conventional dendritic cell’, ‘CD103-positive dendritic cell’, ‘CD8_alpha-positive CD11b-negative dendritic cell’, and ‘dendritic cell’ were relabeled as DC cells.

We ran Tangram2 and Tangram on the paired mouse colon data with default parameters. To calculate AUPRC and AUROC, we took the intersection of genes between the cell culture and colon scRNA-seq datasets, treating the ground truth (GT) genes as positives and the others as negatives. Genes were ranked by logfoldchange and compared against the ground truth.

We compare our results with highly variable genes (HVG), ranked by their dispersion, calculated across the DCs. Highly variable genes are extracted in the same way as for the Tangram2 mapping.

#### B-cell activation

In our analysis of B-cell activation genes, we used one Visium section of human lymph node data (dataset: lymph vis) and unpaired scRNA-seq data from the same tissue (dataset: lymph sc), focusing on Donor A16 as a representative example. We applied the same filtering as in the Slide-tags analysis, excluding mouse-specific features (starting with “MT-”, “MT”, “RP”, and “RPS”) and followed the same normalization and highly variable gene extraction procedures. Cell types were labeled using the “Subset_Broad” annotations. Tangram were run with default settings.

To assess B-cell activation detection between B and T cells, we evaluated three pathways: 1) GO:0042113 – B-cell activation (positive control), 2) GO:0042110 – T cell activation (negative control), and 3) GO:0012501 – programmed cell death (negative control). Pathways were obtained from Gene Ontology (filters: “taxon_subset_closure_label: Homo sapiens”, “type: protein”, and “annotation_class_list_label: {PATHWAY NAME}”, downloaded on 09/20/2024).

We extracted interaction coefficients for the interaction B_naive ← T_CD4+_TfH and ranked genes by their absolute coefficient values, comparing them with pathway membership (positive if present in the pathway, negative otherwise). To establish a baseline for comparison, we subsetted the data to B_naive cells and calculated normalized dispersions for all genes using Scanpy’s highly variable genes method (same as described above). For each ranking (interaction coefficients and normalized dispersions), we calculated the AUROC to assess performance. The AUROC compares true positive rates against false positive rates across varying thresholds, providing a single metric that reflects the ability of our method to prioritize pathway-associated genes.

#### Performance evaluation in Synthetic Data

We applied the synthetic data generation process described earlier to create datasets with various combinations of design parameters: effect_direction {up, down}, signal_effect_scaling {1.5, 2, 4, 5}, signal_effect_base {0.5, 0.8, 0.95, 0.99}, n_spots {100, 250, 500}, and effect_size {10, 25, 50}. These parameters represent:

- effect_direction: whether downstream genes are up- or downregulated,
- signal_effect_scaling: the magnitude of expression changes in interacting cells,
- signal_effect_base: the average expression level of non-interacting cells (quantile-based),
- n_spots: the number of spatial locations,
- effect_size: the number of affected genes.

We generated three replicates per parameter combination using different random seeds, resulting in 864 datasets (864 = 2 * 4 * 4 * 3 * 3 * 3). Preprocessing of the scRNA-seq data followed the same procedure as the Slide-tags analysis. For each dataset, we ran Tangram2 and Tangram2-CCC with default settings, calculating AUROC based on interaction coefficients, treating the artificially introduced effect genes as positives and all others as negatives.

#### Tangram2 analysis of TNBC and ER+ dataset

The breast cancer dataset included scRNA-seq and Visium data from two cancer subtypes, TNBC and ER+. We applied Tangram2-mapping to the scRNA-seq and Visium dataset from the same cancer subtype. Data normalization is first applied to the scRNA-seq, and then the top 100 marker genes from each cell type are combined and used as the training gene for Tangram2-mapping. Tangram2-mapping integrated mode is applied here. **Supplementary Figure 5A** shows the pathology annotation of one TNBC patient, while **Supplementary 5B** gives the predicted distribution of all nine major cell types. The mapping result of major cell types is consistent with the histopathological annotation as well as prior result by Stereoscope^39^: the normal epithelial is mainly mapped to the normal breast tissue region while the cancer epithelial mainly stays in the region of ductal carcinoma in-situ (DCIS) or invasive cancer; T cells are localized in the lymphocyte aggregates; the CAFs and macrophages are localized in spatial proximity with cancer epithelial cells, in the invasive cancer region. The B cells tend to have sparse and confined mapping, highly colocalized with other immune cells such as T cells. Similar patterns have been observed in previous studies and have been correlated to clinical outcome ^57^. After mapping, the colocalization relationship between cell types is calculated by obtaining the Pearson coefficient between the cell type composition at spot level. The colocalization heatmap (**Figure 4A**) is obtained by averaging the Pearson coefficient from each patient of the same cancer subtype. The cell types with overall compositions less than 0.05% are removed from the analysis.

Tangram2-CCC is then applied to the TNBC dataset to study the immunosuppressive mechanisms in the tumor microenvironment. After obtaining the interaction coefficients, we extract and visualize the predefined gene list from the relevant cell types.

#### Macrophage-Treg interaction and TSK analysis in cSCC

The cSCC dataset included scRNA-seq and Visium data from six patients (P2, P4, P5, P6, P9, P10). Two adjacent Visium sections were obtained for P4 and P6, and three for P2, P5, P9, and P10. We excluded genes beginning with (mt, mir, ac0, linc, itgb, rps, rpl, rp) and removed all isoforms (with numeric suffixes) as well as genes containing “orf” or “-as.” Data normalization followed the methods described earlier. Analyses were conducted per patient by mapping each patient’s scRNA-seq data to their combined Visium sections. Interaction coefficients were averaged across patients for all analyses.

We mainly examined Mac ← Treg interaction. Top genes in the interaction were identified using the Kneedle algorithm (kneed.KneeLocator(ranks,interaction_coefficient,S=5, curve=“convex”, direction=“decreasing”) from kneed v.0.8.5).In the pathway analysis we utilized gp.enrichr from the gseapy package (v.1.1.3) with gene_sets=‘MSigDB_Hallmark_2020’. For survival analysis, we used TCGA datasets (TCGA_lusc, TCGA_hnsc, TCGA_skcm) with z-score normalized RNA-seq data. Enrichment of the Mac ← Treg signature was scored via scanpy.tl.score_genes. Patients were stratified into high and low quartiles based on scores, with 1st and 4th quartiles representing low and high classes, respectively. Kaplan-Meier curves were generated with kaplan_meier_estimator (sksurv v.0.23.1) using “OS_STATUS” for event and “OS_MONTHS” for time, with log-rank tests to calculate p-values.

Colocalization analysis of cSCC samples was performed using the same approach as that applied to breast cancer patients. The Pearson correlation coefficient was used to quantify the degree of colocalization between cell types. Cell populations with an overall composition below 0.05% were excluded from the analysis to minimize noise from rare cell types. To investigate the immunosuppressive mechanisms of TSK cells, interaction coefficients were computed and visualized for a predefined list of genes from the relevant cell types.

## Data Availability

No new primary human-subject data were collected. The dataset used to reproduce the figures and results in the paper can be found at: https://zenodo.org/records/17594748 Public datasets used in this study include:

- slide-tags: (https://singlecell.broadinstitute.org/single_cell/study/SCP2169/slide-tags-snrna-seq-on-human-tonsil)
- cell cultures: https://www.ncbi.nlm.nih.gov/geo/query/acc.cgi?acc=GSE135382
- colon: https://cellxgene.cziscience.com/collections/4fa07e63-f712-4d8c-b885-2c515b5e2743
- lymph vis: https://support.10xgenomics.com/spatial-gene-expression/datasets/1.1.0/V1_Human_Lymph_Node?
- lymph sc: https://cell2location.cog.sanger.ac.uk/paper/integrated_lymphoid_organ_scrna/RegressionNBV4Torch_57covariates_73260cells_10237genes/sc.h5ad
- cscc: https://www.ncbi.nlm.nih.gov/geo/query/acc.cgi?acc=GSE144240
- TCGA Lung Squamous Cell Carcinoma (TCGA, PanCancer Atlas): https://www.cbioportal.org/study/summary?id=lusc_tcga_pan_can_atlas_2018
- TCGA Head and Neck Squamous Cell Carcinoma: https://www.cbioportal.org/study/summary?id=hnsc_tcga_pan_can_atlas_2018
- TCGA Skin Cutaneous Melanoma: https://www.cbioportal.org/study/summary?id=skcm_tcga_pan_can_atlas_2018
- breast: https://cellxgene.cziscience.com/e/9fddb063-056d-4202-8b8a-4b0ee531d3ce.cxg/ (donor_id: CID4465).

## Code Availability

The Tangram2 package as well as the code to reproduce the result in the manuscript can be found at https://github.com/Genentech/tangram2

## Author Contribution

These authors contributed equally: Hejin Huang, Alma Andersson.

H.H, A.A, A.B. and T.B. conceived and designed the study. H.H developed the Tangram2-mapping module and performed the mapping and deconvolution benchmarks. A.A developed the Tangram2-CCC and Tangram2-evalkit modules and validated CCC. H.H analyzed TNBC datasets, and H.H. and A.A jointly analyzed the cSCC datasets. S.Z.W, J.C and S.M. guided the biological data analysis. K.H.H. contributed to the data-benchmarking pipeline. J.-C. H.. helped design the loss function of Tangram2-mapping. S.G. assisted with code implementation. A.B. and T.B. provided supervision and general guidance throughout the project. G.S., G.H., H.CB., S.T., S.M. and D.R. provided mentorship and high-level feedback. H.H. and A.A. wrote the manuscript.

## Acknowledgement

We thank L. Gaffney for her help with figure preparation and H.V. Assel for valuable discussions regarding the optimization functions of Tangram2-mapping. Figure templates from BioRender.com were used in their creation.

## Supplementary Figures

**Supplementary Figure 1.**
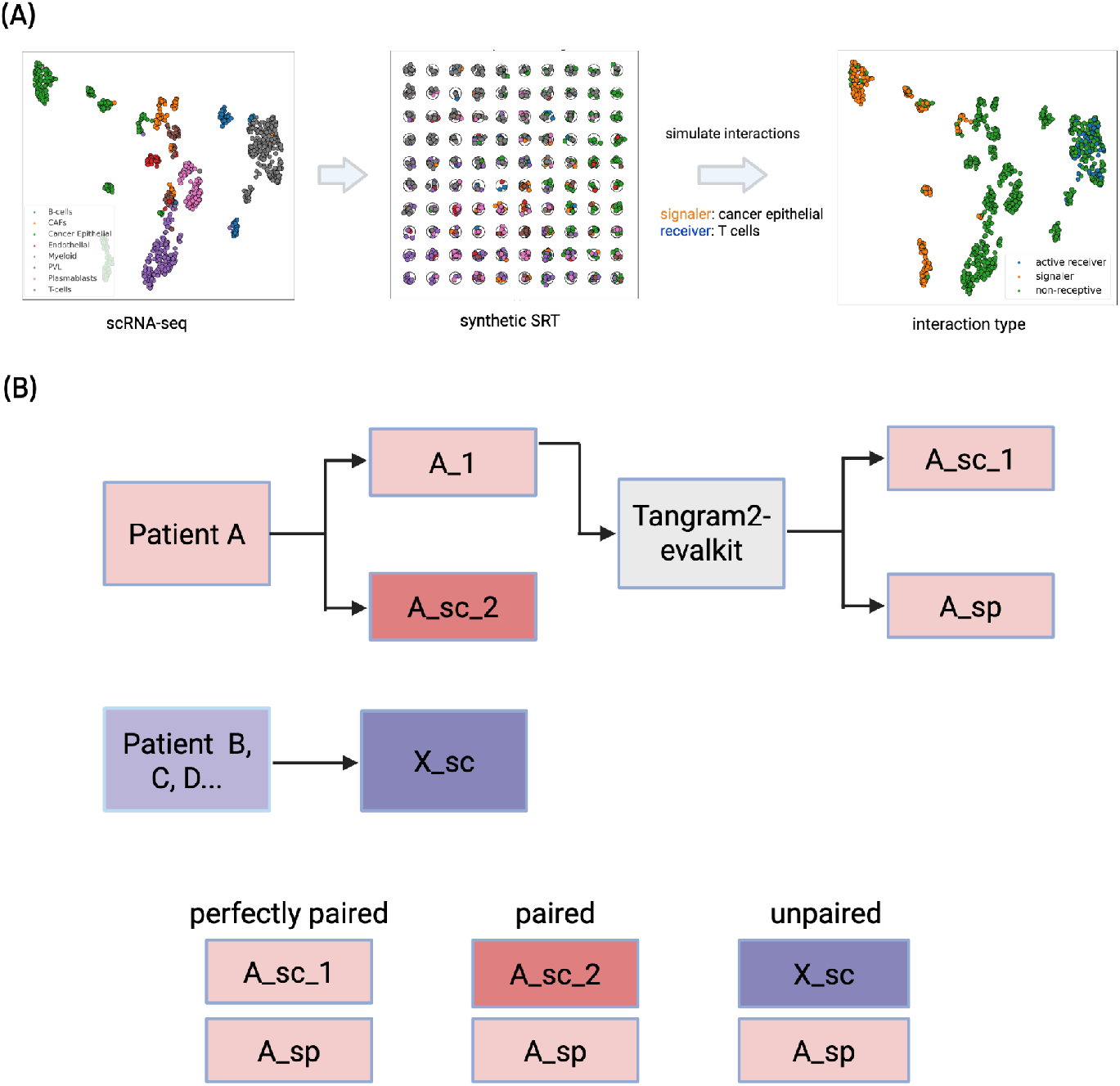
**A**. Generation of simulated datasets for deconvolution and mapping evaluation **B**. data generation under three different scenarios: when the SRT and scRNA-seq are perfectly paired, paired and unpaired. The paired and unpaired datasets are used for Tangram2-mapping evaluations.

**Supplementary Figure 2.**
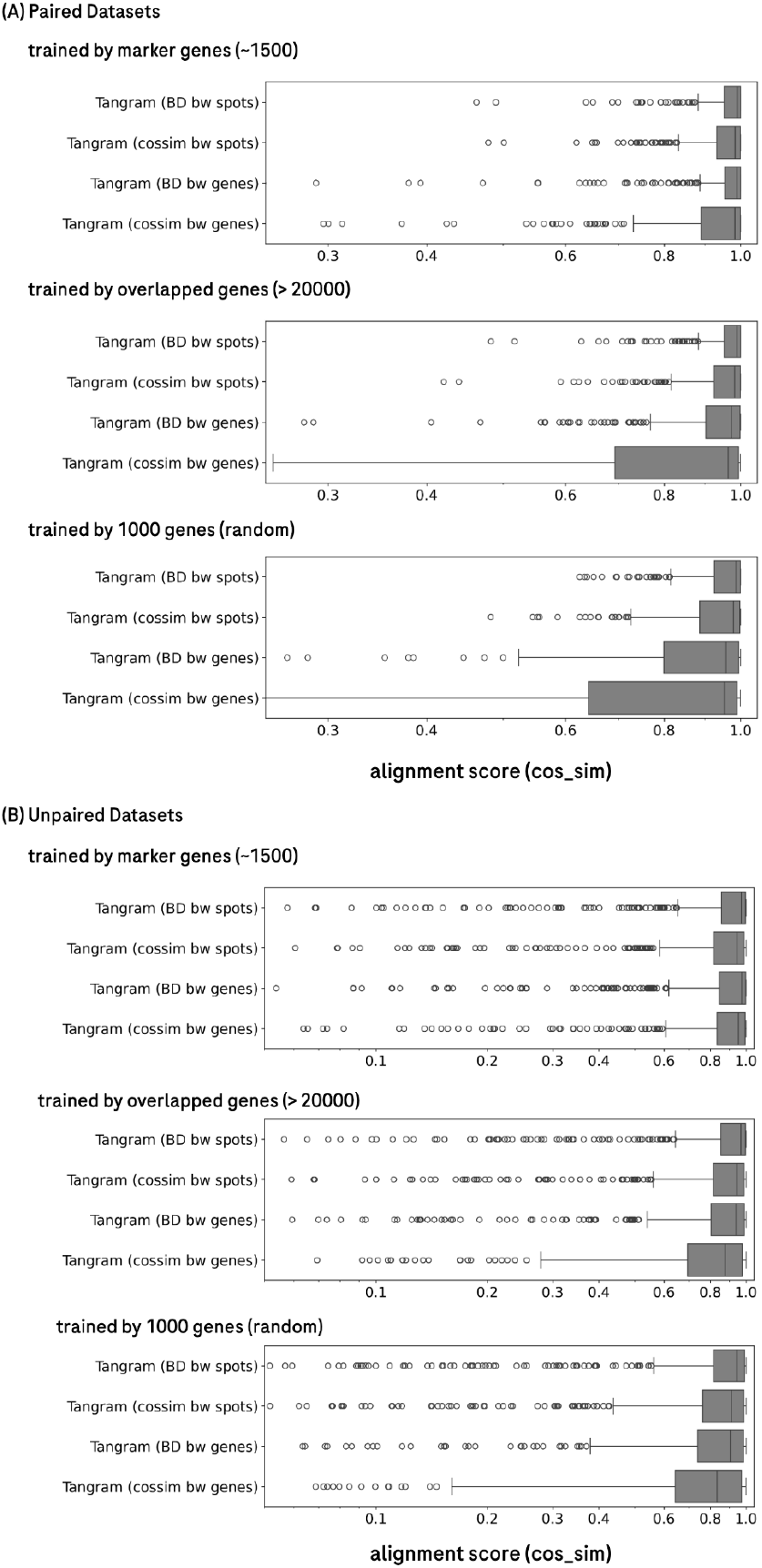
Ablation study shows that the Bhattacharyya distance between spots gives the highest and most robust performance for both paired and unpaired data.

**Supplementary Figure 3.**
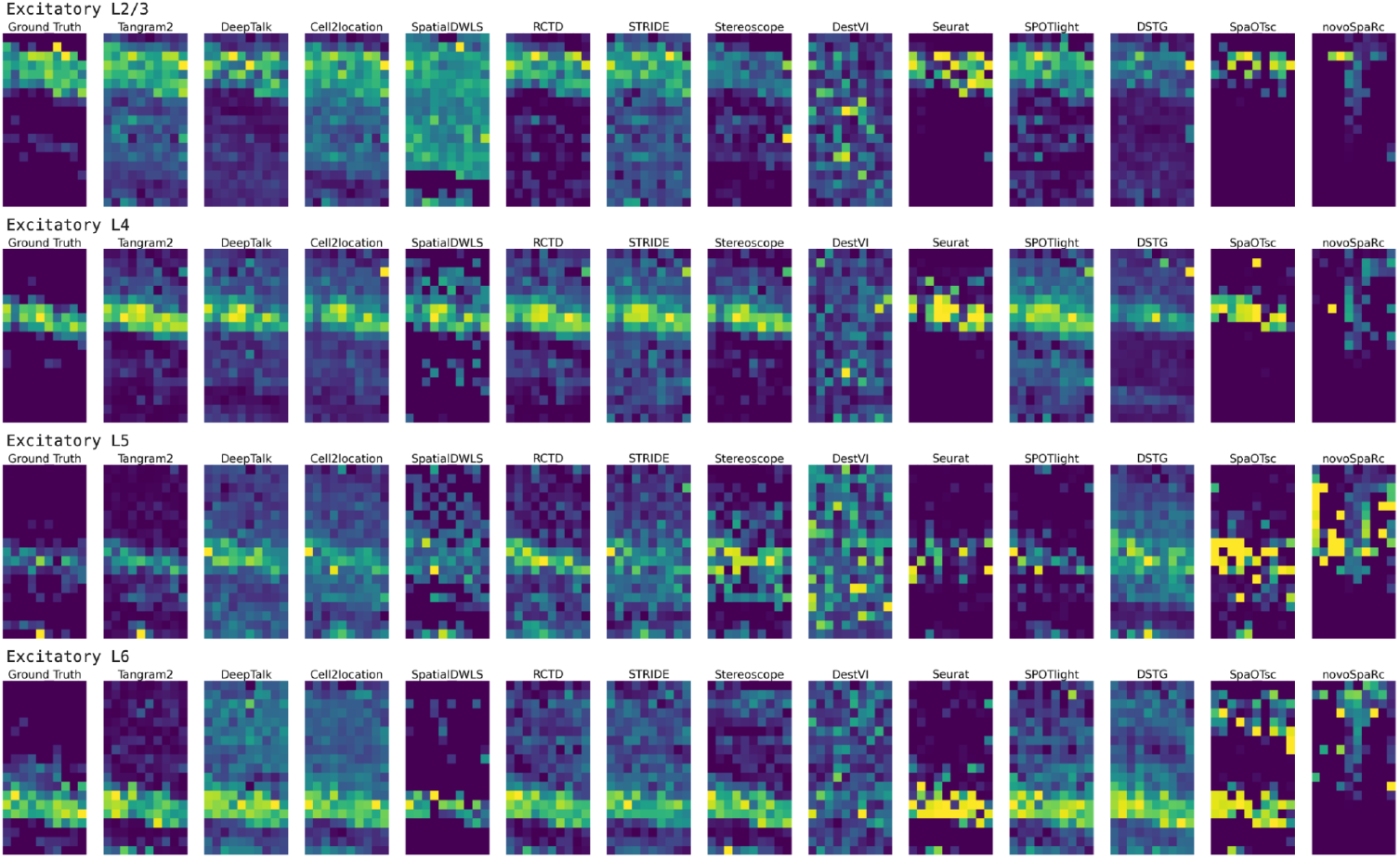
Tangram2-mapping gives the prediction closest to ground truth in the pseudobulked STARMAP datasets. The predicted excitatory neuron distribution of all public tools compared with ground truth from pseudo-bulked STARMAP data from Li et al.

**Supplementary Figure 4.**
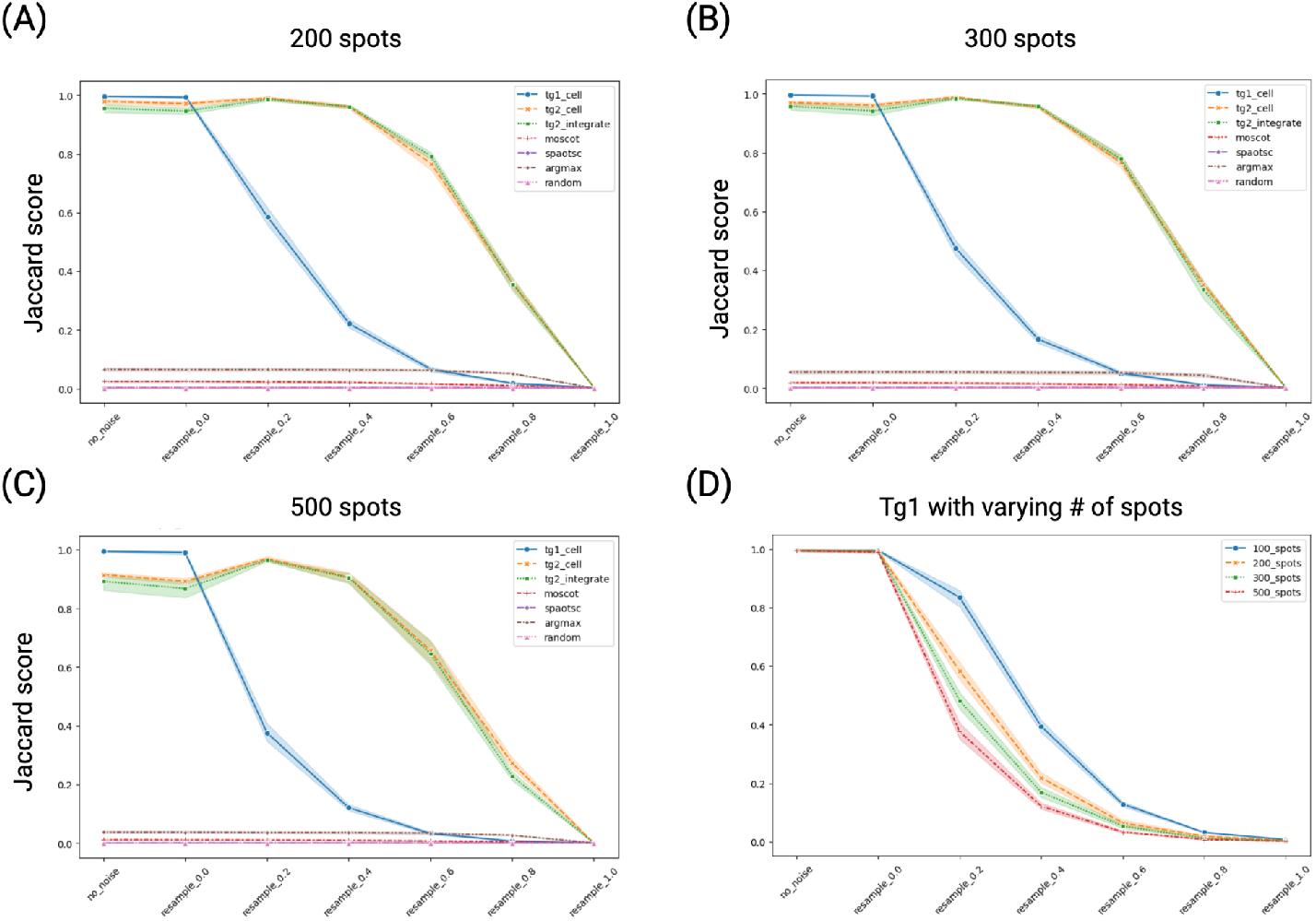
Tangram2-mapping gives state-of-art mapping accuracy across varying numbers of spots. Additional mapping accuracy evaluation result: the comparison of Jaccard scores across all mapping tools at different noise levels when the total number of spots is (A) 200, (B) 300 and (C) 500; (D) the performance of Tangram2 with respect to the varying number of spots from 100 to 500.

**Supplementary Figure 5.**
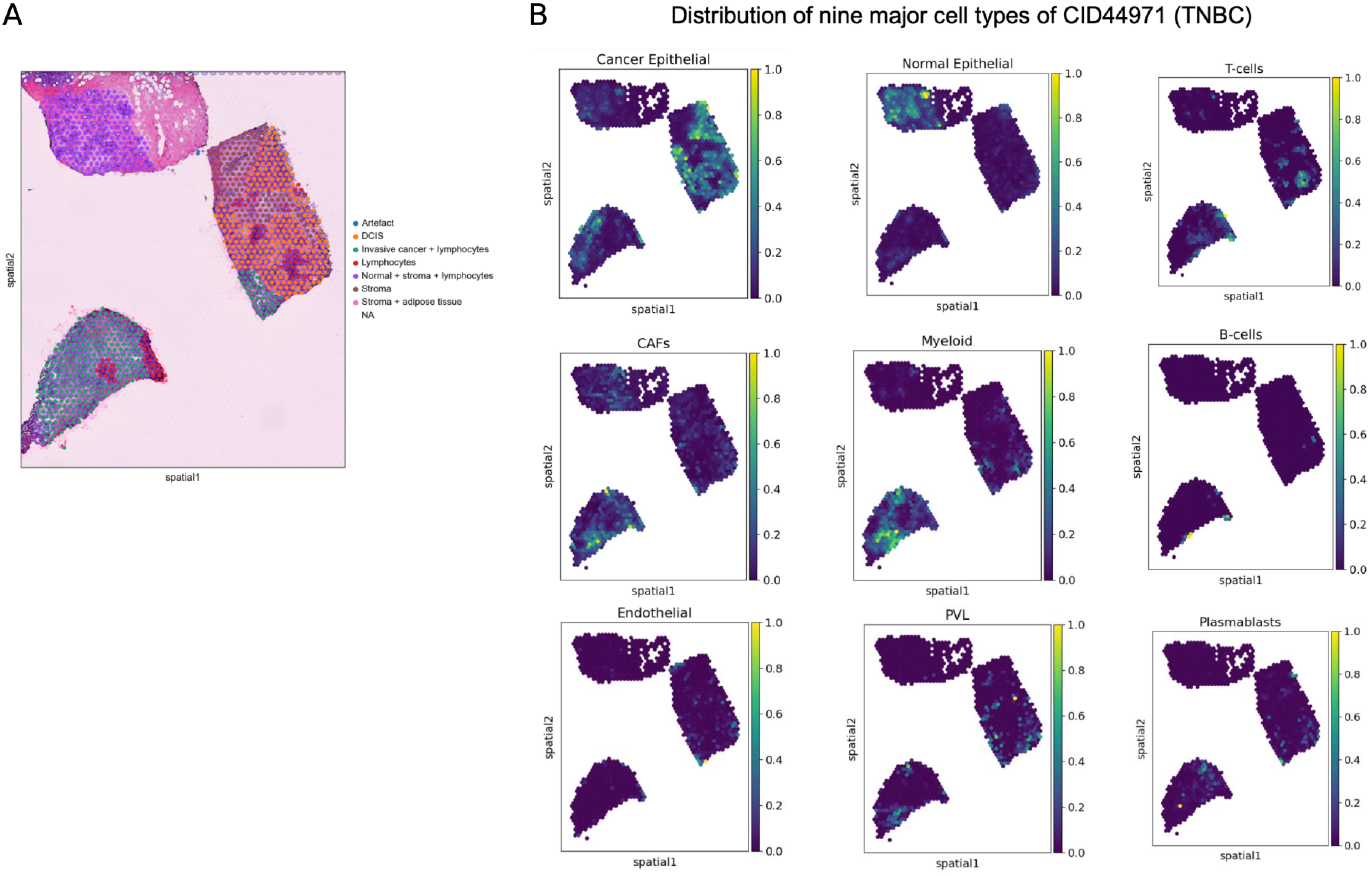
Tangram2-mapping enables accurate alignment between scRNA-seq and spatial transcriptomics. A) the pathologist’s annotation of the Visium sample from one TNBC patient; B) Tangram2-mapping predicted the distribution of nine major cell types of on patient (CID44971).

**Supplementary Figure 6.**
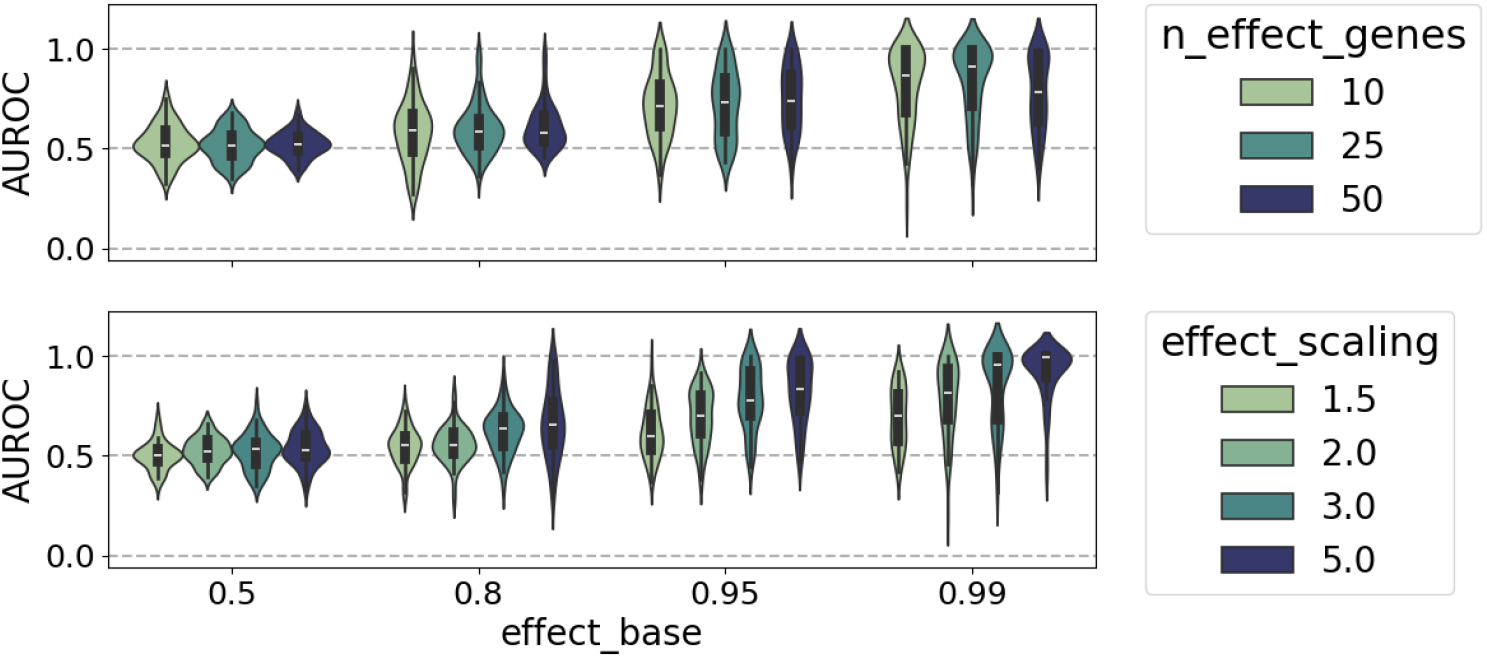
AUROC of Interaction predictions across varying synthetic conditions. Performance on the synthetic data with artificial interactions when stratified by two different axes effect_base (baseline expression) and either of n_effect_genes, and effect_scaling.

**Supplementary Figure 7.**
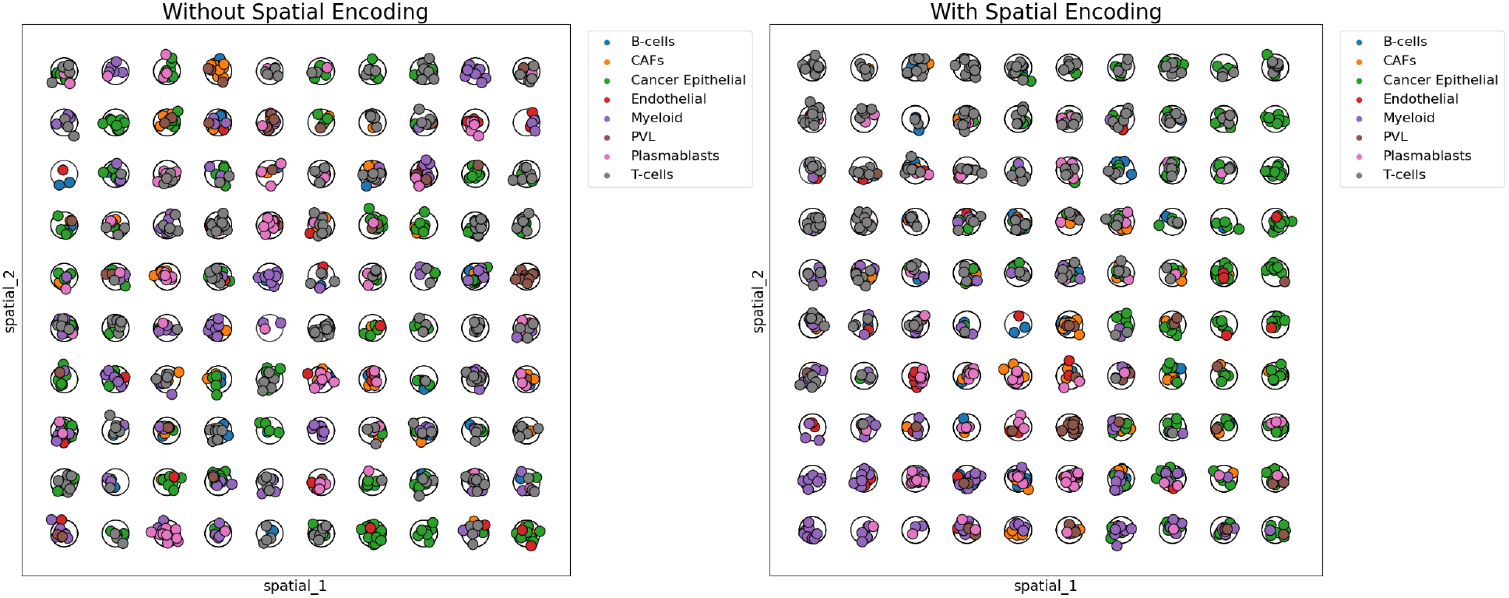
Spatial encoding creates a spatially structured distribution of cell types. Same synthetic dataset with (right) and without (left) spatial encoding enabled. The only difference between the two datasets is the spatial arrangement of the spots. In the spatial encoding case, proximal spots have more similar cell type compositions.

**Supplementary Figure 8.**
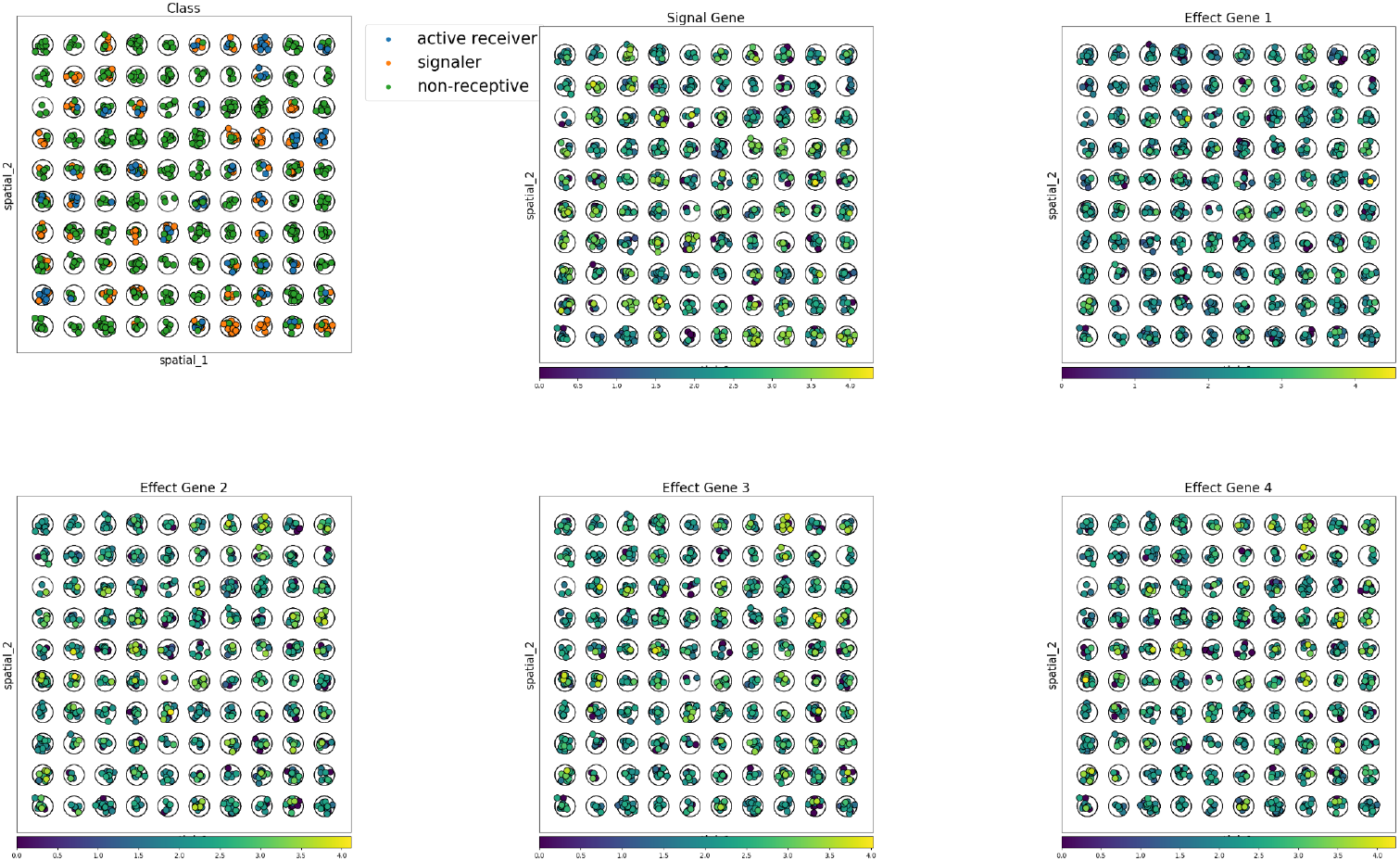
The expression of effect genes align with the spatial distribution of signaling and receiving cells. Spatial plots of the different classes (top left), signal gene expression (top middle), and a sample of the effect genes (rest). Cells are colored by their class or gene expression levels. In this dataset only Cancer Epithelial can be senders and only T-cells were receptive. In total 10 effect genes were added.

**Supplementary Figure 9.**
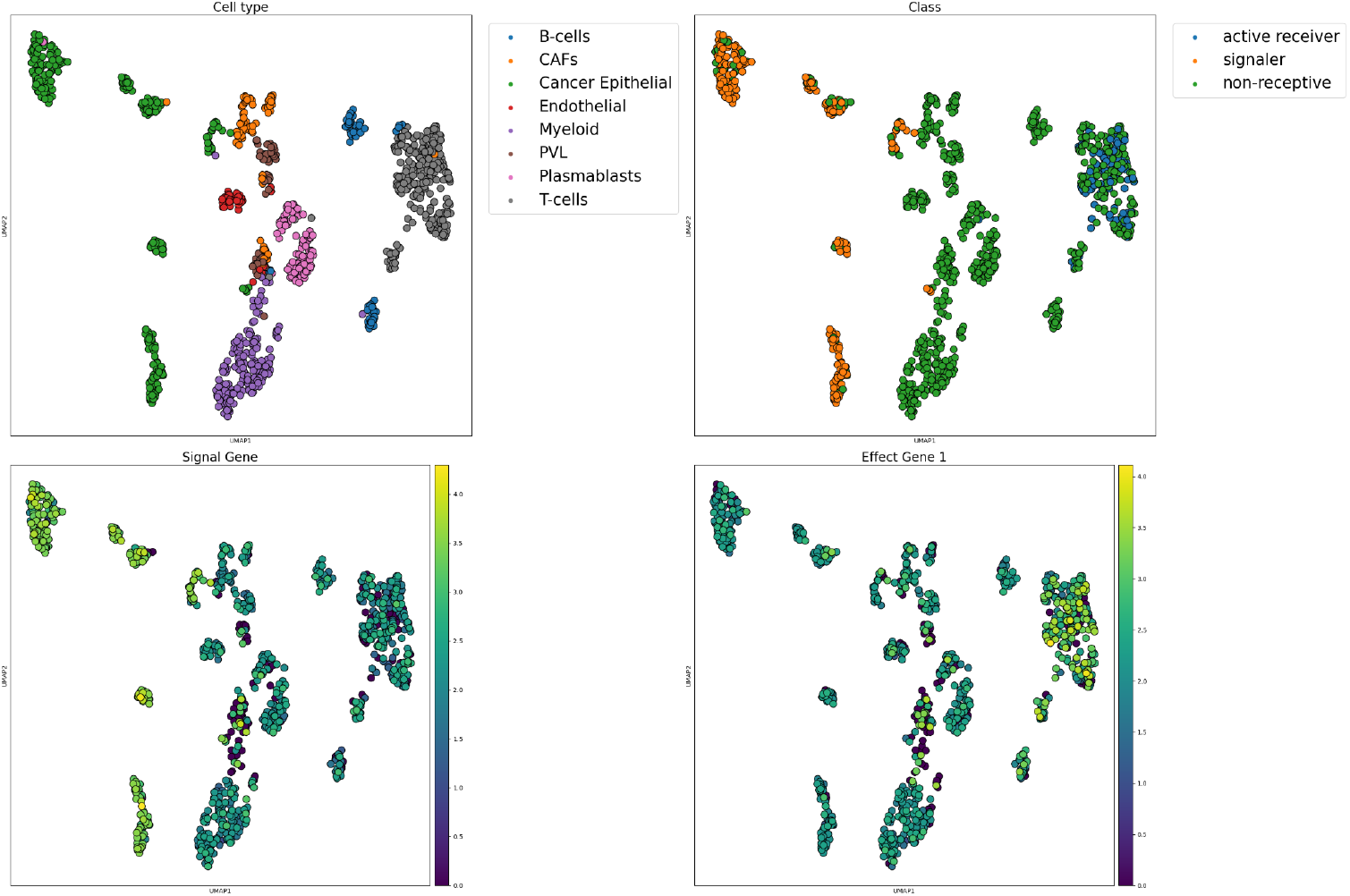
Adding artificial interactions does not impact cell states. UMAP visualization of the synthetic dataset shown in Supplementary Figure 8. The subplots show: (top left) – cell type, (top right) – class, (bottom left) the signal gene expression, (bottom right) – expression of one of the effect genes (effect 1). The cell type identity still dominates the variance in the data and the signal/effect genes do not seem to perturb the gene expression significantly.

**Supplementary Figure 10.**
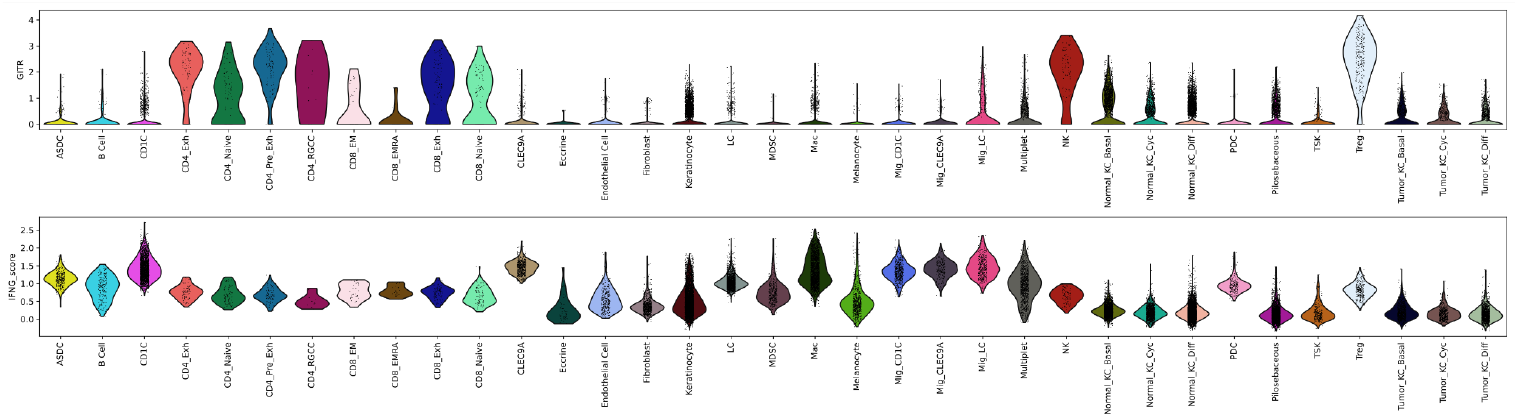
Comparisons of GITR expression (up) and INFγ score (bottom) among all cell types. Treg shows the highest GITR expression and relatively high INFγ score.

**Supplementary Figure 11.**
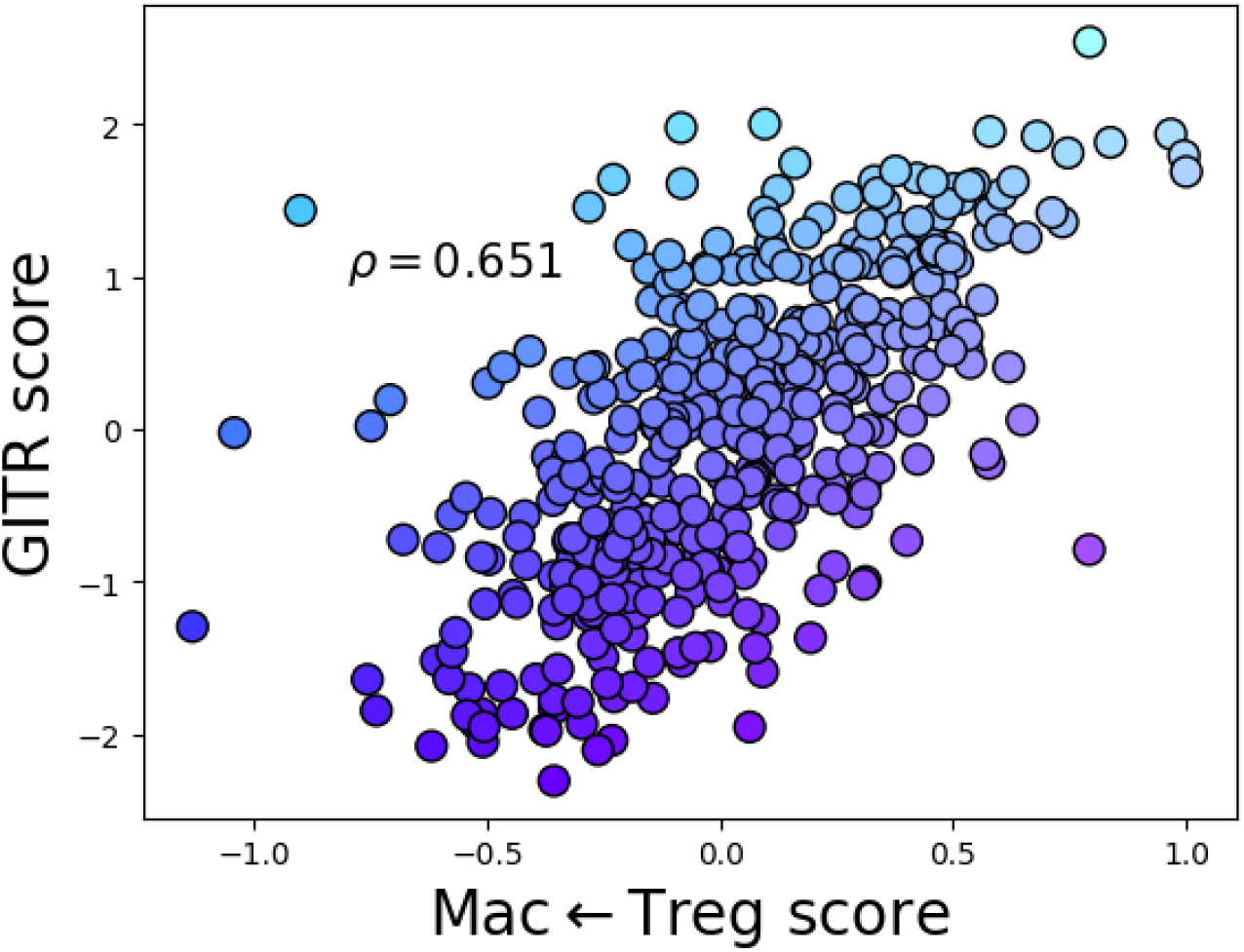
GITR scores and Macrophage ←Treg scores correlate. Each dot is a patient, x-axis shows the patient’s Macrophage ← Treg score and the y-axis the GITR score. The ρ-value is the Spearman correlation (p-value: p = 8.8 × 10^-53^).

**Supplementary Figure 12.**
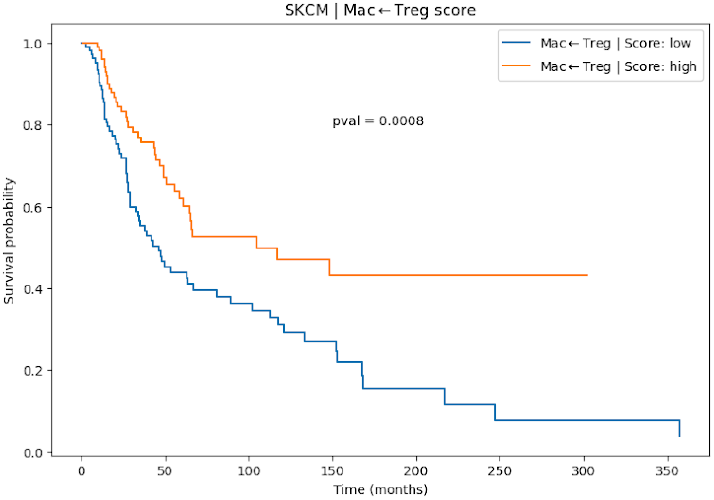
Macrophage ← Treg score stratifies patients w.r.t. clinical outcome. Kaplan-Meyer plot between the top and bottom quartile of SKCM patients based on their Macrophage ← Treg signature score.

